# A set of orthogonal versatile interacting peptide tags for imaging cellular proteins

**DOI:** 10.1101/2022.12.14.520515

**Authors:** Alexa Suyama, Kaylyn L. Devlin, Miguel Macias-Contreras, Julia K. Doh, Ujwal Shinde, Kimberly E. Beatty

## Abstract

Genetic tags are transformative tools for investigating the function, localization, and interactions of cellular proteins. Most studies today are reliant on selective labeling of more than one protein to obtain comprehensive information on a protein’s behavior in situ. Some proteins can be analyzed by fusion to protein tag, such as green fluorescent protein, HaloTag, or SNAP-Tag. Other proteins benefit from labeling via small peptide tags, such as the recently reported versatile interacting peptide (VIP) tags. VIP tags enable observations of protein localization and trafficking with bright fluorophores or nanoparticles. Here we expand the VIP toolkit by presenting two new tags: TinyVIPER and PunyVIPER. These two tags were designed for use with MiniVIPER for labeling up to three distinct proteins at once in living cells. Labeling is mediated by the formation of a high affinity, biocompatible heterodimeric coiled coil. Each tag was validated by fluorescence microscopy, including observation of transferrin receptor 1 trafficking in live cells. We verified that labeling via each tag is highly specific, with no cross-reactivity between the three VIP tags under cellular conditions. Lastly, the self-sorting tags were used for simultaneous labeling of three protein targets (i.e., TOMM20, histone 2B, and actin), highlighting their utility for multicolor microscopy. MiniVIPER, TinyVIPER, and PunyVIPER are small and robust peptide tags for selective labeling of cellular proteins.

## Introduction

Imaging by fluorescence microscopy (FM) is a powerful method for investigating how cellular proteins organize in space and time. Such studies typically use fluorescent labels to visualize selected proteins or cellular features. Organelles are often labeled using small molecule fluorescent stains that function in relation to the organelle’s properties, such as membrane potential (mitochondria), DNA intercalation (nucleus), or pH (lysosomes)^1^. However, the shortage of protein-specific stains restricts them primarily to organelle imaging^2^.

In comparison, protein targets are usually identified by immunolabeling or fusion to a genetic tag. Immunolabeling has several major drawbacks. First, many antibodies lack specificity, resulting in dubious observations and poor reproducibility^3^. A 2008 study found that 49% of antibodies failed to produce a staining pattern consistent with prior work or bioinformatics data^4^. Second, immunolabeling is typically done post-fixation, precluding dynamic studies. Third, immunolabeling protocols can introduce cell damage and artifacts, again leading to misinformation^5^.

An excellent alternative to immunolabeling are genetic tags, where fluorescent labeling is achieved by the fusion of a protein tag to the protein of interest. There are many such tags for imaging in living cells^6^. The most prominent example is the green fluorescent protein (GFP)^7^. Although there are many differently colored fluorescent proteins (FPs), they all have spectral properties determined (and limited) by a post-translationally generated chromophore^8^. Two enzymes have also found widespread use as protein tags: a haloalkane dehalogenase (HaloTag)^9^ and a DNA alkyltransferase (SNAP-tag)^10^. Enzyme tags accept fluorescent substrate mimics, which covalently bind within their active site. These tags confer a significant advantage over FPs since their substrates can be synthesized to exhibit optimal fluorescent properties (e.g., quantum yield, emission, photostability, etc.). However, these and other protein tags^11^ are relatively large (14-30 kDa), which can disrupt folding, trafficking, binding, or function of the target protein^12^.

Peptide tags (<8 kDa) are less likely to impact the tagged protein’s physiological structure or function. An early example was the tetracysteine tag (<1 kDa)^13^, although issues with toxicity and background labeling have limited its widespread use. In contrast, coiled coils have emerged as a versatile and biocompatible motif for labeling cellular proteins^14, 15^. They are comprised of a short peptide tag (4.3 – 7 kDa) that forms an alpha-helical coiled coil heterodimer with a labeled peptide (“probe peptide”). Probe peptides are live cell membrane impermeant, which is ideal for labeling and observing dynamic receptors. The specificity of coiled coils is determined by their peptide sequence, with interstrand salt bridges and a hydrophobic interface driving dimerization-mediated labeling^16^. By separating the tag from the label, coiled coils can maintain high labeling specificity while minimizing the impact on the target. Another advantage is the ability to easily change reporters using variably labeled probe peptides^17^.

The first example of the coiled-coil tag approach was reported by Matsuzaki and coworkers^18^, who used an E3-K3 heterodimer^19^ to label membrane receptors for imaging. Since 2008, several other groups have used coiled-coil tags for cell imaging^15^. Tamamura and coworkers have developed fluorogenic tags^20, 21^, including one for imaging within living cells^22^. Some have developed probe peptides that mediate a proximity-induced reaction to ligate a fluorophore to a receptor^21, 23^. One approach (“Peptide-PAINT”) used transient coiled coil binding for super-resolution microscopy^24^.

Since 2017, we have described three coiled-coil tags termed versatile interacting peptide (VIP) tags. VIP tags are small (4.3 to 6.2 kDa) and can be used for various imaging applications^17, 25, 26-28^. In 2017, we reported our first tag for live cell imaging: VIP Y/Z^26^. Next, we described VIPER, a tag formed by dimerization between a genetically-encoded tag (Coil<u>E</u>) and a probe peptide (Coil<u>R</u>)^28^. We showed that VIPER could highlight organelles or track receptor endocytosis in live cells by “pulse-chase” labeling or time-lapse imaging. Live, intracellular labeling was achieved through light-mediated release of probe peptides from hollow gold nanoshells^27^. VIPER also illustrated that VIP tags enable labeling with reporters matched to the application, such as quantum dots (Qdots) for correlative light and electron microscopy (CLEM)^28^. Our CLEM studies included quantitative analysis of receptor labeling, by which we found that VIPER labeling was more efficient than immunolabeling. Most recently, we developed the MiniVIPER tag^17^, which was inspired by a coiled-coil heterodimer first reported by Vinson and coworkers^29^. MiniVIPER is comprised of a MiniE and MiniR heterodimer. We showed that MiniVIPER could be used with either VIPER or VIP Y/Z for labeling two cellular proteins at once^17^. However, those tags were not designed to be orthogonal to MiniVIPER, limiting the utility of VIP tags for multitarget labeling.

As prior studies suggested it would be feasible to create a set of orthogonal coiled-coil tags^30, 31^, the goal of our current work was to design a set of self-sorting peptide tags optimized for multiprotein labeling in cells. We aimed for this set of peptides to be biocompatible and bioorthogonal for stable, high-affinity labeling. To this end, we created two new tags, TinyVIPER and PunyVIPER, which were designed de novo to be used with MiniVIPER. We characterized each tag in vitro and found that the coiled-coil heterodimers interacted with low nanomolar affinity and high stability. In cells, each tag enabled selective labeling of proteins, including transferrin receptor 1 (TfR1), actin, TOMM20, and histone 2B (H2B). We show that VIP-tagged TfR1 retained physiological ligand binding and internalization patterns. Pairwise and three-way combinations of these tags showed bright, target-specific labeling, without cross-reactivity, of up to three distinct proteins in cells.

## Materials and Experimental Details

See the Electronic Supporting Information (ESI) for a detailed description of all materials and methods used.

## Results and Discussion

### Design of new self-sorting VIP tags

In the current work, we present a set of six peptides that self-sort to form coiled-coil heterodimeric tags for protein labeling without cross-reactivity (**Figure 1**). We de novo designed TinyVIPER and PunyVIPER to be orthogonal to our previously reported MiniVIPER tag^17^. Each tag is 4.3 kDa, making them among the smallest tags reported to date. Peptide tag sequences, properties, and structural depictions (i.e., helical wheel diagrams) are reported in **Figure 2**.

**Figure 1.**
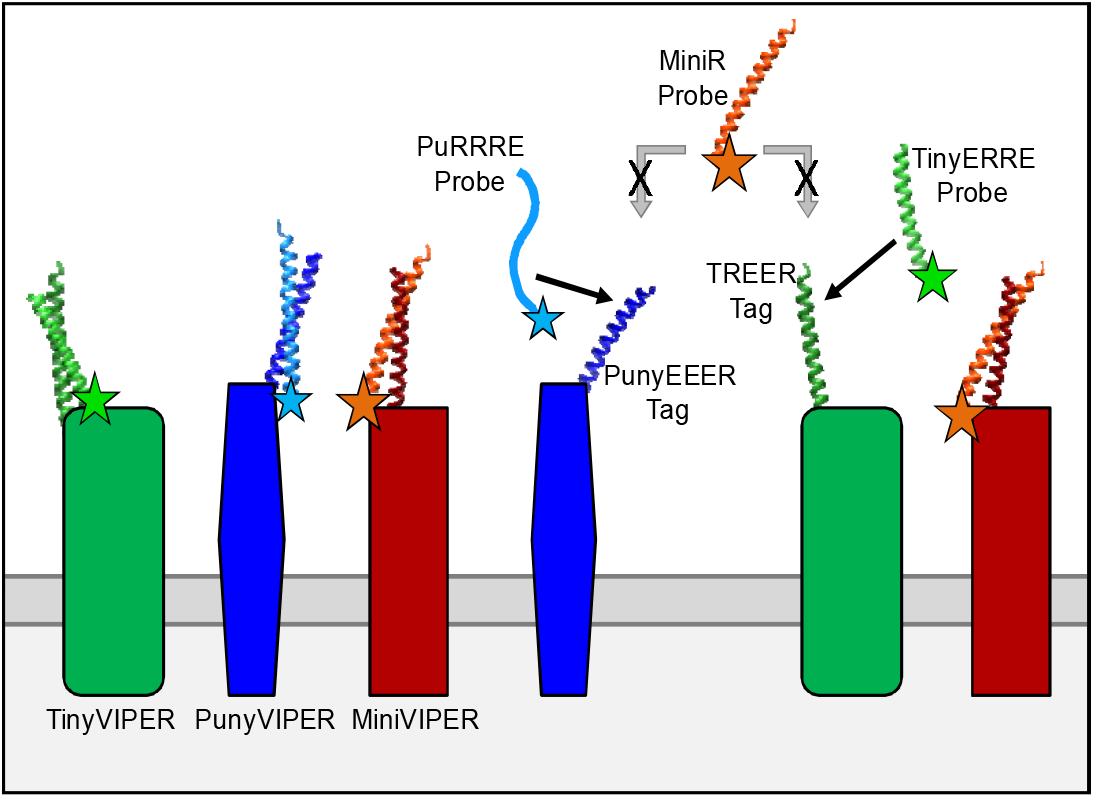
Cellular protein labeling via orthogonal, self-sorting VIP tags. A set of three peptide tags, MiniVIPER (red), TinyVIPER (green), and PunyVIPER (blue), enable selective protein labeling by formation of coiled-coil heterodimers between each tag and a corresponding fluorescent probe peptide.

**Figure 2.**
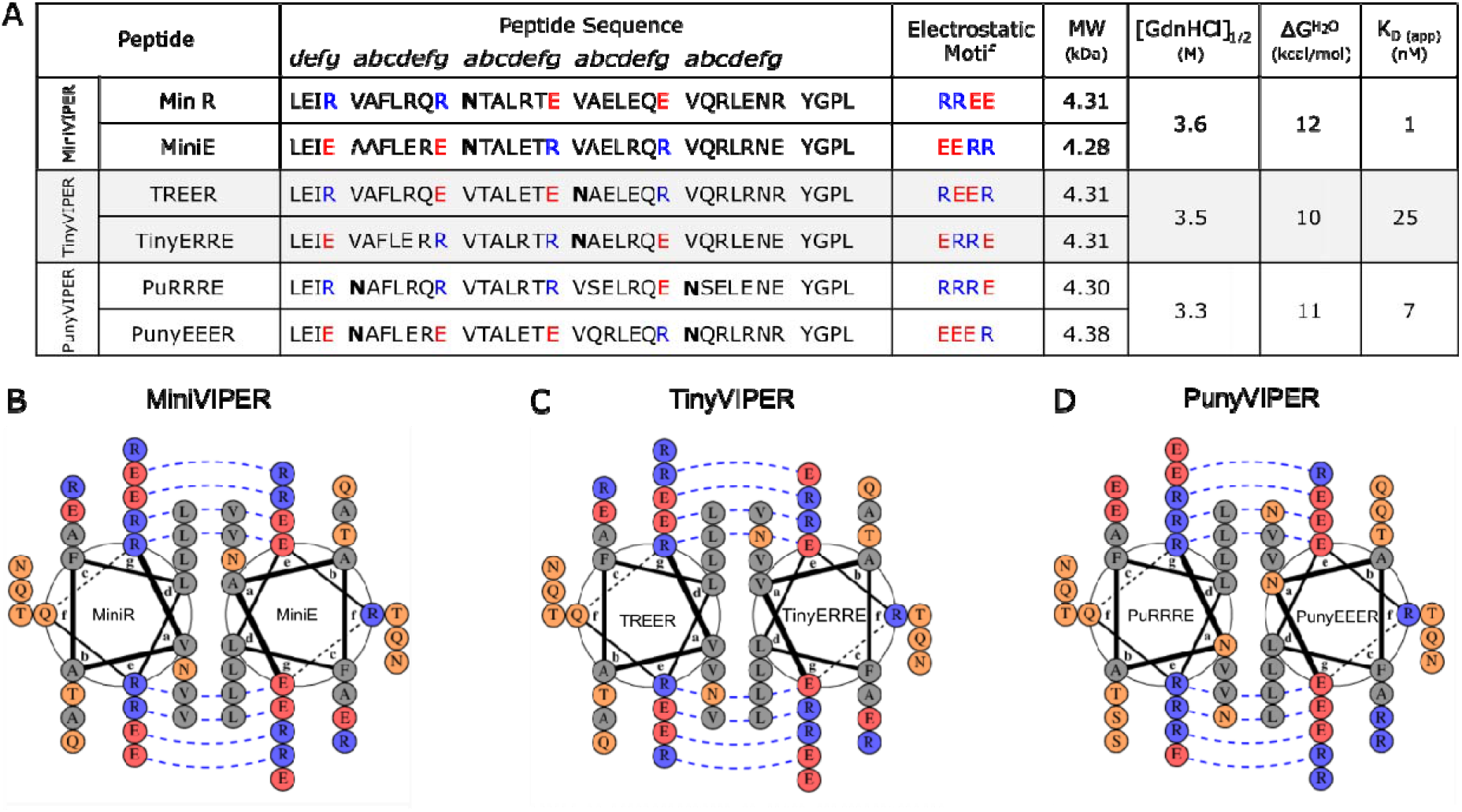
Design and biophysical properties of orthogonal VIP tags. (A) Peptide sequences reported using single-letter amino acid code, displayed relative to heptad motif in header. Positively- and negatively-charged amino acids that define the electrostatic motif are labeled in blue and red, respectively. Asn (N) matches are bolded. Biophysical properties were determined by circular dichroism spectroscopy. The denaturant GdnHCl concentration where the coiled coil is 50% folded is reported as [GdnHCl]_1/2_. The free energy of heterodimer formation without denaturant (ΔG^H2O^) was calculated by linear extrapolation and the dissociation constant, K_D (app)_, was calculated from the ΔG^H2O^. (B)-(D) Helical wheel diagrams for each heterodimeric coiled-coil. Residues are color-coded: charged = blue (+) or red (-); polar = orange; hydrophobic = grey. Favorable salt bridges are illustrated by a blue dashed line. Diagrams were generated using DrawCoil1.0^38^ (https://grigoryanlab.org/drawcoil/).

Each dimeric pair has characteristic alpha-helical coiled coil interactions, including heptad repeats (denoted *abcdefg*), salt bridges, and a hydrophobic interface. Each tag is four heptads, with charge distribution designed to ensure interaction specificity and orthogonality. Residues at the *a* and *d* positions form a hydrophobic interface between the two coils that drives heterodimer formation, while interstrand salt bridges (*g*↔*e*′) enforce heterodimer specificity^32^. We used a Glu-Arg electrostatic pair, which is 0.35 kcal/mol more stable than the Glu-Lys pair that is commonly used in other coiled-coil dimers^30, 33^. There are eight favorable salt bridges upon the formation of each optimized parallel heterodimer: MiniVIPER (**Figure 2B**), TinyVIPER (**Figure 2C**), or PunyVIPER (**Figure 2D**). TinyVIPER is a product of T<u>REER</u>–Tiny<u>ERRE</u> heterodimer formation, with the charge distribution indicated in the peptide name. PunyVIPER is a heterodimer of Pu<u>RRRE</u> and Puny<u>EEER</u>.

Alpha helical coiled coils can form parallel and anti-parallel dimers. We included an *a* position Asn-Asn match to promote formation of parallel dimers by hydrogen bonding between opposing coils^30, 34-36^. The Asn-Asn match was included in the second heptad of MiniVIPER, the third heptad for TinyVIPER, and the first and fourth for PunyVIPER. The remaining *a* position residues were Val, which further enhances binding specificity because Val-Asn interactions are disfavored^35^. Various potential homodimers and mismatched coiled-coil pairs are not expected to form due to destabilizing interactions^36, 37^, as illustrated in **Figure S1** (see **ESI**).

### Generation of recombinant peptides for analysis and imaging

Each peptide was made by recombinant expression in *E. coli*. Peptide sequences included a coil followed by a short linker, reactive Cys, and C-terminal hexahistidine tag (for purification). The Cys was included to enable site-specific attachment of a fluorophore. In prior work, we made probe peptides using standard thiol-maleimide conjugation^28^ (i.e., Tris buffer pH 7.2 with excess TCEP and reactive fluorophore) or Weiss’s solid-state labeling method^17, 39^. However, ligation was often inefficient. Unmodified and fluorophore-labeled peptides could not be separated, resulting in labeling reagents contaminated with up to 70% non-fluorescent peptides^17^. For the current work, we optimized labeling by adapting a protocol published by Watts and coworkers^40^. This method quenches excess reducing agent (i.e., TCEP) before the addition of excess reactive fluorophore. After adopting this quenching step, we routinely achieved 50-90% labeling efficiencies (**Table S4**). We encountered a surprising difficulty attaching a fluorophore to TREER. Therefore, TREER was used solely as the genetic tag for TinyVIPER labeling. Our method of making probe peptides uses commercially available fluorophores available in a range of colors and properties, making it feasible for others to rapidly generate probe peptides.

### Biophysical characterization of coiled-coil heterodimers

The secondary structure of the peptides and heterodimers were analyzed by circular dichroism (CD) spectroscopy. The ellipticity of each peptide alone was measured between 200-260 nm. All peptides, except for PuRRRE, had a distinct curve with minima at 208 nm and 222 nm indicative of alpha-helical structure^41^ (**Figure 3A**). In contrast, PuRRRE was unstructured. We cannot rule out homodimer formation at the high peptide concentrations required for these measurements, but the electrostatics would make such interactions disfavored (see Figure S1). Heterodimers were evaluated as equimolar mixtures of two peptides (i.e., MiniE + MiniR, PunyEEER + PuRRRE, or TinyERRE + TREER). The CD spectra were consistent with the formation of alpha-helical heterodimers (**Figure 3B** and **Figure S2**).

**Figure 3.**
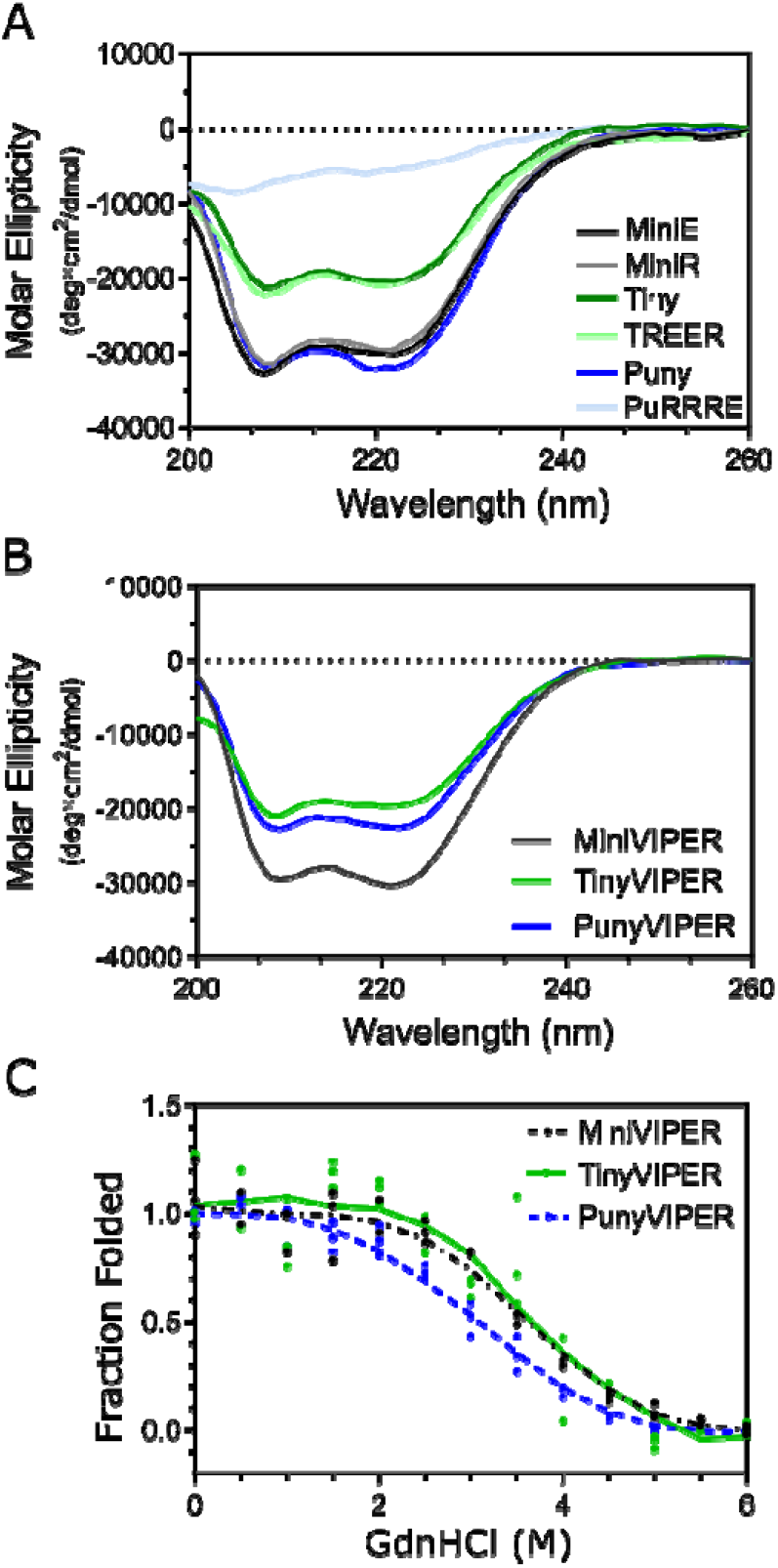
MiniVIPER, TinyVIPER, and PunyVIPER form stable alpha helical heterodimers. CD spectra of peptide monomers (A) and coiled-coil dimers (B). (C) Denaturation curves for MiniVIPER, TinyVIPER, and PunyVIPER. In (B) and (C), each peptide was present at a 1:1 molar ratio with 10 μM total peptide (5 μM each). Each graph represents three experimental replicates.

Next, we analyzed the conformational stability of heterodimers in increasing concentration of the denaturant guanidine hydrochloride (GdnHCl; 0 to 6 M) by measuring ellipticity at 222 nm. **Figure 3C** shows the fraction of heterodimer folded, assuming that 100% was folded below 1 M GdnHCl and 0% was folded at 6 M GdnHCl. The denaturation curve suggested that all three dimers had high stability, with each remaining 50% folded at ∼3.5 M GdnHCl. The observed ellipticity of each dimer with denaturant was used to extrapolate the free energy of unfolding in absence of denaturant (ΔG^H2O^). All three heterodimers were found to have similar ΔG^H2O^ (∼11 kcal/mol) (**Figure 2A**).

The denaturation curves were used to calculate the apparent dissociation constant (K_D (app)_), using the method described by Litowski and Hodges^19^. Each heterodimer had low nanomolar affinity (<25 nM) (**Figure 2A**). These values are consistent with the low nanomolar affinity measured for other four heptad coiled coils^42^ and match our prior prediction for MiniVIPER’s K_D_^17^. The CD measurements required peptides to be at micromolar concentrations in order to be detected, raising the possibility that the true K_D_ is quite a bit lower^43^.

### Receptor trafficking observed with TinyVIPER and PunyVIPER

We established that each new VIP tag enabled specific labeling of proteins in living cells by imaging TfR1, a key component of the iron uptake machinery. We selected TfR1 as a model protein due to its well characterized localization, trafficking, and ligand (transferrin) binding^44^. Chinese hamster ovary (CHO) TRVb cells^45^ expressing tagged receptors were cooled to pause endocytosis before treatment with fluorescent transferrin (Tf-488) and 100 nM Sulfo-Cyanine 5 (Cy5)-labeled probe peptide. Cells were fixed immediately (0 min) or after receptor endocytosis (10 min at 37 °C) and then imaged by confocal FM.

We found that each VIP tag enabled selective labeling of TfR1 (**Figure 4**).

Cells expressing VIP-tagged TfR1 showed distinct signal at the cell surface (0 min) and in endocytic vesicles upon internalization (10 min). Cells expressing untagged TfR1 bound Tf-488, but not probe peptide. Signal from membrane-associated tagged TfR1 colocalized with Tf-488, as measured by Pearson’s correlation coefficient (PCC) (0 min = 0.60-0.83), and colocalization was maintained upon endocytosis (10 min = 0.47-0.63) (**Figure S3**). These colocalization metrics provide strong support that VIP-tagged receptor retained ligand-binding function.

**Figure 4.**
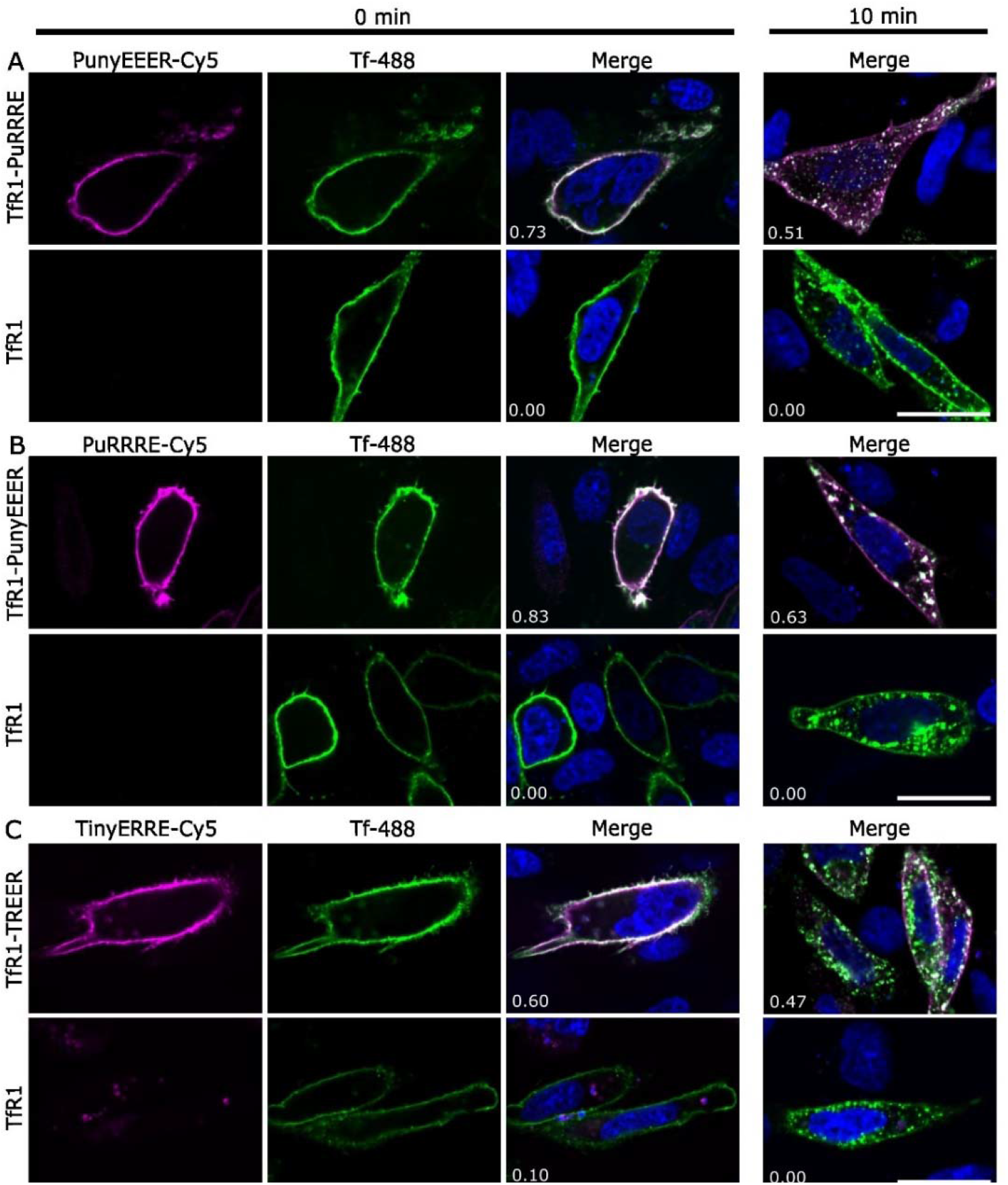
VIP-tagged TfR1 receptor localizes to the membrane, binds Tf, and trafficks by endocytosis. CHO TRVb cells expressing VIP-tagged TfR1 were labeled live (4 °C) with 100 nM Cy5-conjugated probe peptide and 50 ng/mL Tf-488. Cells were fixed immediately after labeling (0 min) or after incubation at 37 °C to allow receptor-ligand internalization (10 min). (A) Cells expressing TfR1-PuRRRE or untagged TfR1. (B) Cells expressing TfR1-PunyEEER or TfR1 labeled with PuRRRE-Cy5 and Tf-488. (C) Cells expressing TfR1-TREER or TfR1 labeled with TinyERRE-Cy5 and Tf-488. Micrographs are false-colored (Cy5, magenta; Tf-488, green; Hoechst, blue) and green-magenta overlap appears white in the merge. The PCC between the Cy5 and 488 channels is reported in the lower left of each merge micrograph. Scale bars represent 20 μm.

We did not observe Cy5 signal in untransfected cells or cells expressing untagged TfR1, illustrating that labeling was highly specific. We occasionally observed probe peptide associated with cellular debris, as shown in the lower left panel in **Figure 4C**.

For PunyVIPER (**Figure 4A, 4B**) and MiniVIPER^17^ (**Figure S4**), we demonstrated that either peptide could be encoded as the tag. We expect the same to be true of TinyVIPER, although we were limited to the use of TREER as the tag in absence of fluorescent TREER probe peptide. All together, these results show that PuRRRE, PunyEEER, TREER, MiniE, and MiniR tags all enabled selective labeling of proteins in live cells.

### VIP-mediated two-color labeling of organelles

After each tag was validated individually, we next used pairs of VIP tags to label two distinct targets in cells. We selected proteins with defined localization patterns, including actin (cytoskeleton), TOMM20 (mitochondria), and histone 2B (H2B; nucleus). First, we showed that TinyVIPER (actin) and PunyVIPER (mitochondria) could be used together to label cellular proteins without cross-reactivity (**Figure 5**). Human osteosarcoma cells (U-2 OS) expressing mEmerald (mEm)-Actin-TREER and TOMM20-PuRRRE were fixed and treated with probe peptides (TinyERRE-Cy3 and PunyEEER-Cy5). PunyVIPER (Cy5) labeling was restricted to mitochondria, which retained normal morphology. TinyVIPER (Cy3) labeled actin, with signal colocalized with green fluorescence (mEmerald), as expected. We did not observe Cy5 signal in the cytoskeleton or Cy3 signal in mitochondria, suggesting that the two tags are orthogonal.

**Figure 5.**
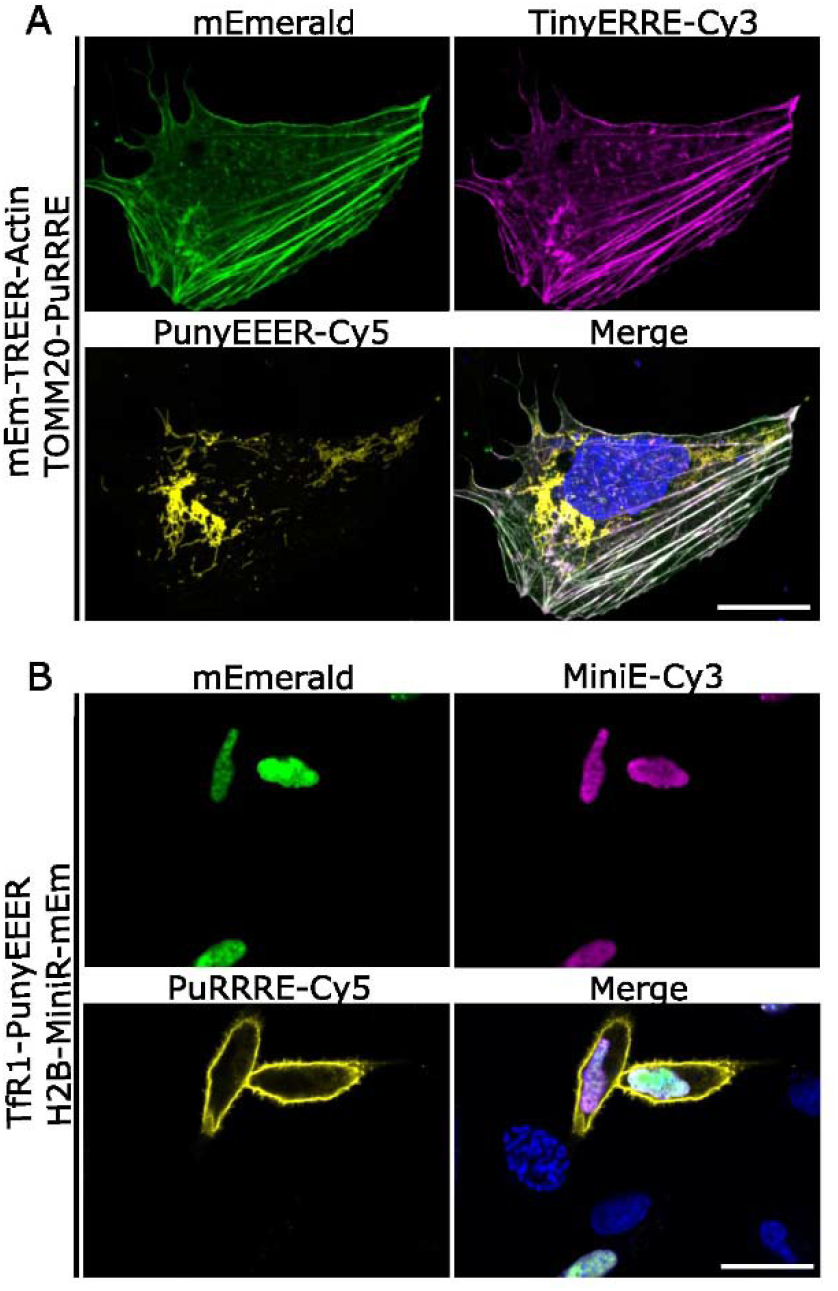
VIP tags enable specific labeling of two protein targets in cells. (A) U-2 OS cells expressing mEm-TREER-Actin and TOMM20-PuRRRE were treated with PunyEEER-Cy5 and TinyERRE-Cy3 to enable simultaneous visualization of mitochondria and actin filaments. Micrographs are presented as a maximum intensity projection of a confocal z-stack. (B) CHO TRVb cells expressing TfR1-PunyEEER and H2B-MiniR-mEm were treated with PuRRRE-Cy5 (live) and MiniE-Cy3 (post-fixation). Micrographs are false-colored (Cy3, magenta; Cy5, yellow; mEm, green) and the merge includes Hoechst nuclear stain (blue). Scale bars represent 20 μm.

Next, we evaluated PunyVIPER and MiniVIPER for imaging H2B (nucleus) and TfR1 (cell membrane) in CHO TRVb cells (**Figure 5B**). Cells expressing TfR1-PunyEEER and H2B-MiniR-mEm were first labeled with PuRRRE-Cy5 live, to restrict labeling to membrane localized receptor. Cells were then fixed, treated with MiniE-Cy3, and imaged by confocal FM. Again, VIP-mediated labeling was specific and tagged proteins retained their physiological organelle localization. MiniE-Cy3 colocalized with mEmerald within the nucleus.

In both dual-labeling experiments, there was no nonspecific labeling in untransfected cells, cells expressing a single VIP-tagged protein, or cells transfected to express untagged proteins (**Figure S5**). Moreover, VIP tags did not appear to affect the normal localization of actin, TOMM20, H2B, or TfR1. Additional combinations of two VIP tags are included in **Figure 6**.

Altogether, these studies illustrate that MiniVIPER, TinyVIPER, and PunyVIPER can be used to label and observe two protein targets simultaneously by FM.

**Figure 6.**
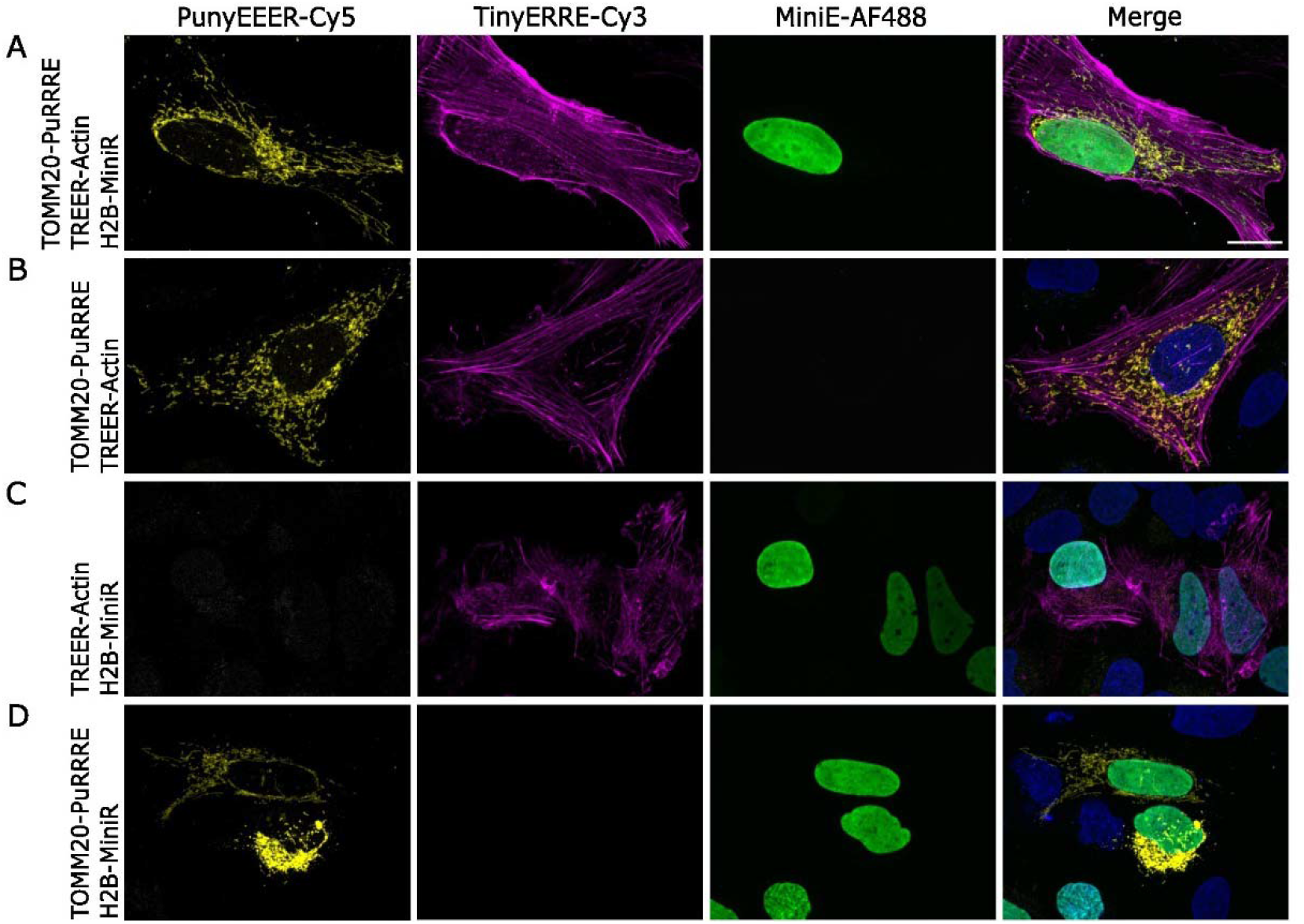
MiniVIPER, TinyVIPER, and PunyVIPER enable simultaneous imaging of three distinct protein targets in cells. (A) U-2 OS cells expressing TREER-Actin, H2B-MiniR, and TOMM20-PuRRRE were fixed and then treated simultaneously with PunyEEER-Cy5, TinyERRE-Cy3, and MiniE-AF488. (B)-(D) Cells expressing two VIP-tagged proteins (as indicated) and labeled with all three probe peptides. Cells were imaged by confocal FM imaging and micrographs represent maximum intensity projections from z-stacks. Micrographs are false-colored (Cy5, yellow; Cy3, magenta; AF488, green; Hoechst, blue). Scale bars represent 20 μm.

### Simultaneous VIP labeling of three distinct cellular targets

Researchers often label multiple cellular proteins for imaging in order to capture the complexity of protein-protein interactions, dynamics, or reorganization under various conditions. There have been numerous discoveries made through multicolor labeling using large genetic tags (e.g., GFP or HaloTag). By comparison, very few multicolor labeling studies have capitalized on the advantageous properties of peptide tags.

Our objective herein was to create a set of bioorthogonal coiled-coil tags that expand the ability to label multiple proteins in cells. Therefore, we lastly investigated whether TinyVIPER, PunyVIPER, and MiniVIPER could be used together to label three distinct cellular targets without cross-reactivity. We chose to label the cytoskeleton (actin), the nucleus (H2B), and mitochondria (TOMM20) due to their distinct sub-cellular localizations. Fixed cells were labeled simultaneously with three spectrally-distinct probe peptides (i.e., PunyEEER-Cy5, TinyERRE-Cy3, and MiniE-AF488) and then imaged (**Figure 6**). We observed selective labeling of each intended protein target by its corresponding probe peptide. Notably, there was no cross-labeling between mismatched pairs nor non-specific labeling in cells (**Figure S6**). These results provide direct evidence that MiniVIPER, TinyVIPER, and PunyVIPER are orthogonal peptide tags that can be used to simultaneously label three cellular targets.

## Conclusion

As reported here, the VIP tag repertoire has been expanded to include TinyVIPER and PunyVIPER for concurrent use with MiniVIPER. These coiled-coil heterodimers have high affinity (K_D(app)_ <25 nM) and stability, even under strongly denaturing conditions (3M [GdnHCl]_1/2_). These tags are ideal for labeling multiple cellular proteins at once due to several key characteristics. Foremost, they form a selective, bioorthogonal set of coiled-coil tags with no observed cross-reactivity. They are small tags (4.3 kDa), making them unlikely to disrupt physiological protein functions, interactions, or localization. Our studies of VIP-tagged TfR1 support this assertion. Additionally, each tag is compatible with observing receptor trafficking in living cells. Lastly, VIP tags can be used to label proteins with diverse reporters, including fluorophores, biotin, and electron dense particles^17, 28^.

These features highlight VIP tags as an advantageous alternative to conventional immunolabeling or large protein tags (e.g., fluorescent proteins). Additionally, these features suggest their utility for other advanced imaging applications. The live cell compatibility of these tags would allow dynamic imaging of multiple cell receptors in parallel. Furthermore, this set of tags has the potential to advance labeling methodologies available for cell-based electron microscopy (EM) and correlative fluorescence and EM, where labeling methods for target identification are currently limited. In future work we plan to use VIP tags for multicolor CLEM studies of receptor organization. Overall, the VIP toolkit, including TinyVIPER, PunyVIPER, and MiniVIPER, are ideal tools for multiprotein investigations through imaging.

## Acknowledgements

The authors are grateful to our colleagues at Oregon Health & Science University for their advice and guidance. Special appreciation is due to S. Kaech Petrie and the staff of the Advanced Light Microscopy Core, S. Reichow, and C. Enns. We appreciate the significant contributions of past lab members to the development of VIP tags.

## Funding

This research was supported by generous funding from Oregon Health & Science University and the National Institutes of Health (R01 GM122854).

## Notes

### Competing Interest Statement

The authors have declared no competing interest.

## References

(1) Xu, W.; Zeng, Z.; Jiang, J. H.; Chang, Y. T.; Yuan, L. Discerning the Chemistry in Individual Organelles with Small-Molecule Fluorescent Probes. Angew Chem Int Ed Engl 2016, 55 (44), 13658–13699. DOI: 10.1002/anie.201510721

(2) Adhikari, S.; Nice, E. C.; Deutsch, E. W.; Lane, L.; Omenn, G. S.; Pennington, S. R.; Paik, Y.-K.; Overall, C. M.; Corrales, F. J.; Cristea, I. M.; et al. A high-stringency blueprint of the human proteome. Nat Commun 2020, 11 (1), 5301–5301. DOI: 10.1038/s41467-020-19045-9 The Molecular Probes Handbook A Guide to Fluorescent Probes and Labeling Technologies; Life Technologies Corporation, 2010.

(3) Uhlen, M.; Bandrowski, A.; Carr, S.; Edwards, A.; Ellenberg, J.; Lundberg, E.; Rimm, D. L.; Rodriguez, H.; Hiltke, T.; Snyder, M.; et al. A proposal for validation of antibodies. Nat. Meth. 2016, 13 (10), 823–827. DOI: 10.1038/nmeth.3995.

(4) Uhlén, M.; Björling, E.; Agaton, C.; Szigyarto, C. A.-K.; Amini, B.; Andersen, E.; Andersson, A.-C.; Angelidou, P.; Asplund, A.; Asplund, C.; et al. A Human Protein Atlas for Normal and Cancer Tissues Based on Antibody Proteomics. Molecular & Cellular Proteomics 2005, 4 (12), 1920–1932. DOI: https://doi.org/10.1074/mcp.M500279-MCP200.

(5) Schnell, U.; Dijk, F.; Sjollema, K. A.; Giepmans, B. N. G. Immunolabeling artifacts and the need for live-cell imaging. Nat. Meth. 2012, 9 (2), 152–158, 10.1038/nmeth.1855. DOI: http://www.nature.com/nmeth/journal/v9/n2/abs/nmeth.1855.html#supplementary-information.

(6) Li, C.; Tebo, A. G.; Gautier, A. Fluorogenic Labeling Strategies for Biological Imaging. International Journal of Molecular Sciences 2017, 18 (7), 1473. Liu, J.; Cui, Z. Fluorescent Labeling of Proteins of Interest in Live Cells: Beyond Fluorescent Proteins. Bioconjugate Chemistry 2020, 31 (6), 1587–1595. DOI: 10.1021/acs.bioconjchem.0c00181.

(7) Chalfie, M.; Tu, Y.; Euskirchen, G.; Ward, W.; Prasher, D. Green fluorescent protein as a marker for gene expression. Science 1994, 263 (5148), 802–805. DOI: 10.1126/science.8303295.

(8) Rodriguez, E. A.; Campbell, R. E.; Lin, J. Y.; Lin, M. Z.; Miyawaki, A.; Palmer, A. E.; Shu, X.; Zhang, J.; Tsien, R. Y. The Growing and Glowing Toolbox of Fluorescent and Photoactive Proteins. Trends Biochem. Sci. 2017, 42 (2), 111–129. DOI: http://doi.org/10.1016/j.tibs.2016.09.010.

(9) Los, G. V.; Encell, L. P.; McDougall, M. G.; Hartzell, D. D.; Karassina, N.; Zimprich, C.; Wood, M. G.; Learish, R.; Ohana, R. F.; Urh, M.; et al. HaloTag: A Novel Protein Labeling Technology for Cell Imaging and Protein Analysis. ACS Chem. Biol. 2008, 3 (6), 373–382. DOI: 10.1021/cb800025k

(10) Keppler, A.; Gendreizig, S.; Gronemeyer, T.; Pick, H.; Vogel, H.; Johnsson, K. A general method for the covalent labeling of fusion proteins with small molecules in vivo. Nature Biotech. 2003, 21 (1), 86–89.

(11) Plamont, M.-A.; Billon-Denis, E.; Maurin, S.; Gauron, C.; Pimenta, F. M.; Specht, C. G.; Shi, J.; Quérard, J.; Pan, B.; Rossignol, J.; et al. Small fluorescence-activating and absorption-shifting tag for tunable protein imaging in vivo. Proceedings of the National Academy of Sciences 2016, 113 (3), 497–502. DOI: doi:10.1073/pnas.1513094113. Miller, L. W.; Sable, J.; Goelet, P.; Sheetz, M. P.; Cornish, V. W. Methotrexate conjugates: a molecular in vivo protein tag. Angew Chem Int Ed Engl 2004, 43 (13), 1672–1675. DOI: 10.1002/anie.200352852

(12) Costantini, L. M.; Snapp, E. L. Fluorescent proteins in cellular organelles: serious pitfalls and some solutions. DNA Cell Biol 2013, 32 (11), 622–627. DOI: 10.1089/dna.2013.2172 From NLM.

(13) Griffin, B. A.; Adams, S. R.; Tsien, R. Y. Specific covalent labeling of recombinant protein molecules inside live cells. Science 1998, 281 (5374), 269–272.

(14) Lotze, J.; Reinhardt, U.; Seitz, O.; Beck-Sickinger, A. G. Peptide-tags for site-specific protein labelling in vitro and in vivo. Mol Biosyst 2016, 12 (6), 1731–1745. DOI: 10.1039/c6mb00023a

(15) Yano, Y.; Matsuzaki, K. Live-cell imaging of membrane proteins by a coiled-coil labeling method-Principles and applications. Biochim Biophys Acta Biomembr 2019, 1861 (5), 1011–1017. DOI: 10.1016/j.bbamem.2019.02.009

(16) Mason, J. M.; Arndt, K. M. Coiled coil domains: stability, specificity, and biological implications. Chembiochem 2004, 5 (2), 170–176. DOI: 10.1002/cbic.200300781

(17) Doh, J. K.; Tobin, S. J.; Beatty, K. E. MiniVIPER Is a Peptide Tag for Imaging and Translocating Proteins in Cells. Biochemistry 2020, 59 (33), 3051–3059. DOI: 10.1021/acs.biochem.0c00526.

(18) Yano, Y.; Yano, A.; Oishi, S.; Sugimoto, Y.; Tsujimoto, G.; Fujii, N.; Matsuzaki, K. Coiled-Coil Tag-Probe System for Quick Labeling of Membrane Receptors in Living Cells. ACS Chem. Biol. 2008, 3 (6), 341–345.

(19) Litowski, J. R.; Hodges, R. S. Designing heterodimeric two-stranded alpha-helical coiled-coils. Effects of hydrophobicity and alpha-helical propensity on protein folding, stability, and specificity. J Biol Chem 2002, 277 (40), 37272–37279. DOI: 10.1074/jbc.M204257200

(20) Tsutsumi, H.; Nomura, W.; Abe, S.; Mino, T.; Masuda, A.; Ohashi, N.; Tanaka, T.; Ohba, K.; Yamamoto, N.; Akiyoshi, K.; et al. Fluorogenically Active Leucine Zipper Peptides as Tag–Probe Pairs for Protein Imaging in Living Cells. Angew. Chem. Int. Ed. 2009, 48 (48), 9164–9166. DOI: 10.1002/anie.200903183 Tsutsumi□, H.; Abe□, S.; Mino□, T.; Nomura, W.; Tamamura, H. Intense Blue Fluorescence in a Leucine Zipper Assembly. ChemBioChem 2011, 12 (5), 691–694. DOI: 10.1002/cbic.201000692.

(21) Nomura, W.; Mino, T.; Narumi, T.; Ohashi, N.; Masuda, A.; Hashimoto, C.; Tsutsumi, H.; Tamamura, H. Development of crosslink-type tag-probe pairs for fluorescent imaging of proteins. Peptide Science 2010, 94 (6), 843–852. DOI: 10.1002/bip.21444.

(22) Nomura, W.; Ohashi, N.; Mori, A.; Tamamura, H. An in-cell fluorogenic Tag-probe system for protein dynamics imaging enabled by cell-penetrating peptides. Bioconjug Chem 2015, 26 (6), 1080–1085. DOI: 10.1021/acs.bioconjchem.5b00131

(23) Wang, J.; Yu, Y.; Xia, J. Short Peptide Tag for Covalent Protein Labeling Based on Coiled Coils. Bioconjugate Chem. 2014, 25 (1), 178–187. DOI: 10.1021/bc400498p Reinhardt, U.; Lotze, J.; Zernia, S.; Mörl, K.; Beck-Sickinger, A. G.; Seitz, O. Peptide-Templated Acyl Transfer: A Chemical Method for the Labeling of Membrane Proteins on Live Cells. Angew. Chem. Int. Ed. 2014, 10237–10241. DOI: 10.1002/anie.201403214 Reinhardt, U.; Lotze, J.; Mörl, K.; Beck-Sickinger, A. G.; Seitz, O. Rapid Covalent Fluorescence Labeling of Membrane Proteins on Live Cells via Coiled-Coil Templated Acyl Transfer. Bioconjugate Chemistry 2015, 26 (10), 2106–2117. DOI: 10.1021/acs.bioconjchem.5b00387 Lotze, J.; Wolf, P.; Reinhardt, U.; Seitz, O.; Mörl, K.; Beck-Sickinger, A. G. Time-Resolved Tracking of Separately Internalized Neuropeptide Y2 Receptors by Two-Color Pulse-Chase. ACS Chem. Biol. 2018. DOI: 10.1021/acschembio.7b00999.

(24) Eklund, A. S.; Ganji, M.; Gavins, G.; Seitz, O.; Jungmann, R. Peptide-PAINT Super-Resolution Imaging Using Transient Coiled Coil Interactions. Nano Letters 2020, 20 (9), 6732–6737. DOI: 10.1021/acs.nanolett.0c02620.

(25) Doh, J. K.; Tobin, S. J.; Beatty, K. E. Generation of CoilR Probe Peptides for VIPER-labeling of Cellular Proteins. Bio-protocol 2019, 9 (21), e3412. DOI: 10.21769/BioProtoc.3412. Doh, J. K.; Enns, C. A.; Beatty, K. E. Implementing VIPER for Imaging Cellular Proteins by Fluorescence Microscopy. Bio-protocol 2019, 9 (21), e3413. DOI: 10.21769/BioProtoc.3413 Doh, J. K.; Chang, Y. H.; Enns, C. A.; López, C. S.; Beatty, K. E. Imaging VIPER-labeled Cellular Proteins by Correlative Light and Electron Microscopy. Bio-protocol 2019, 9 (21), e3414. DOI: 10.21769/BioProtoc.3414.

(26) Zane, H. K.; Doh, J. K.; Enns, C. A.; Beatty, K. E. Versatile interacting peptide (VIP) tags for labeling proteins with bright chemical reporters. ChemBioChem 2017, 18 (5), 470–474.

(27) Morgan, E.; Doh, J.; Beatty, K.; Reich, N. VIPERnano: Improved Live Cell Intracellular Protein Tracking. ACS Applied Materials & Interfaces 2019, 11 (40), 36383–36390. DOI: 10.1021/acsami.9b12679.

(28) Doh, J. K.; White, J. D.; Zane, H. K.; Chang, Y. H.; Lopez, C. S.; Enns, C. A.; Beatty, K. E. VIPER is a genetically encoded peptide tag for fluorescence and electron microscopy. Proc Natl Acad Sci U S A 2018, 115 (51), 12961–12966. DOI: 10.1073/pnas.1808626115

(29) Moll, J. R.; Ruvinov, S. B.; Pastan, I.; Vinson, C. Designed heterodimerizing leucine zippers with a ranger of pIs and stabilities up to 10−15 M. Protein Science 2001, 10 (3), 649–655. DOI: https://doi.org/10.1110/ps.39401.

(30) Lebar, T.; Lainscek, D.; Merljak, E.; Aupic, J.; Jerala, R. A tunable orthogonal coiled-coil interaction toolbox for engineering mammalian cells. Nat Chem Biol 2020, 16 (5), 513–519. DOI: 10.1038/s41589-019-0443-y

(31) Reinke, A. W.; Grant, R. A.; Keating, A. E. A Synthetic Coiled-Coil Interactome Provides Heterospecific Modules for Molecular Engineering. J. Am. Chem. Soc. 2010, 132 (17), 6025–6031. DOI: 10.1021/ja907617a. Thompson, K. E.; Bashor, C. J.; Lim, W. A.; Keating, A. E. SYNZIP protein interaction toolbox: in vitro and in vivo specifications of heterospecific coiled-coil interaction domains. ACS Synth Biol 2012, 1 (4), 118–129. DOI: 10.1021/sb200015u

(32) Burkhard, P.; Ivaninskii, S.; Lustig, A. Improving Coiled-coil Stability by Optimizing Ionic Interactions. Journal of Molecular Biology 2002, 318 (3), 901–910. DOI: https://doi.org/10.1016/S0022-2836(02)00114-6.

(33) Krylov, D.; Mikhailenko, I.; Vinson, C. A thermodynamic scale for leucine zipper stability and dimerization specificity: e and g interhelical interactions. The EMBO Journal 1994, 13 (12), 2849–2861. DOI: https://doi.org/10.1002/j.1460-2075.1994.tb06579.x. Plaper, T.; Aupič, J.; Dekleva, P.; Lapenta, F.; Keber, M. M.; Jerala, R.; Benčina, M. Coiled-coil heterodimers with increased stability for cellular regulation and sensing SARS-CoV-2 spike protein-mediated cell fusion. Scientific Reports 2021, 11 (1), 9136. DOI: 10.1038/s41598-021-88315-3.

(34) Gonzalez, L.; Woolfson, D. N.; Alber, T. Buried polar residues and structural specificity in the GCN4 leucine zipper. Nat. Struct. Mol. Biol. 1996, 3 (12), 1011–1018. DOI: 10.1038/nsb1296-1011.

(35) Acharya, A.; Ruvinov, S. B.; Gal, J.; Moll, J. R.; Vinson, C. A heterodimerizing leucine zipper coiled coil system for examining the specificity of a position interactions: amino acids I, V, L, N, A, and K. Biochemistry 2002, 41 (48), 14122–14131. DOI: 10.1021/bi020486r

(36) Zeng, X.; Herndon, A. M.; Hu, J. C. Buried asparagines determine the dimerization specificities of leucine zipper mutants. Proc Natl Acad Sci U S A 1997, 94 (8), 3673–3678. DOI: 10.1073/pnas.94.8.3673

(37) O’Shea, E. K.; Klemm, J. D.; Kim, P. S.; Alber, T. X-ray Structure of the GCN4 Leucine Zipper, a Two-Stranded, Parallel Coiled Coil. Science 1991, 254 (5031), 539–544.

(38) Grigoryan, G.; Keating, A. E. Structural specificity in coiled-coil interactions. Curr Opin Struct Biol 2008, 18 (4), 477–483. DOI: 10.1016/j.sbi.2008.04.008

(39) Kim, Y.; Ho, S. O.; Gassman, N. R.; Korlann, Y.; Landorf, E. V.; Collart, F. R.; Weiss, S. Efficient Site-Specific Labeling of Proteins via Cysteines. Bioconjugate Chem. 2008, 19 (3), 786–791. DOI: 10.1021/bc7002499.

(40) Kantner, T.; Alkhawaja, B.; Watts, A. G. In Situ Quenching of Trialkylphosphine Reducing Agents Using Water-Soluble PEG-Azides Improves Maleimide Conjugation to Proteins. ACS Omega 2017, 2 (9), 5785–5791. DOI: 10.1021/acsomega.7b01094.

(41) Miles, A. J.; Janes, R. W.; Wallace, B. A. Tools and methods for circular dichroism spectroscopy of proteins: a tutorial review. Chem Soc Rev 2021, 50 (15), 8400–8413. DOI: 10.1039/d0cs00558d

(42) De Crescenzo, G.; Litowski, J. R.; Hodges, R. S.; O’Connor-McCourt, M. D. Real-time monitoring of the interactions of two-stranded de novo designed coiled-coils: effect of chain length on the kinetic and thermodynamic constants of binding. Biochemistry 2003, 42 (6), 1754–1763. DOI: 10.1021/bi0268450

(43) Jarmoskaite, I.; Alsadhan, I.; Vaidyanathan, P. P.; Herschlag, D. How to measure and evaluate binding affinities. eLife 2020, 9. DOI: 10.7554/elife.57264.

(44) Zhao, N.; Enns, C. A. Chapter Three - Iron Transport Machinery of Human Cells: Players and Their Interactions. In Current Topics in Membranes, Argüello, J. M., Lutsenko, S. Eds.; Vol. 69; Academic Press, 2012; pp 67–93. Mayle, K. M.; Le, A. M.; Kamei, D. T. The intracellular trafficking pathway of transferrin. Biochimica et Biophysica Acta (BBA) - General Subjects 2012, 1820 (3), 264–281. DOI: 10.1016/j.bbagen.2011.09.009.

(45) McGraw, T. E.; Greenfield, L.; Maxfield, F. R. Functional expression of the human transferrin receptor cDNA in Chinese hamster ovary cells deficient in endogenous transferrin receptor. J Cell Biol 1987, 105 (1), 207–214. DOI: 10.1083/jcb.105.1.207

