## Supporting Information for "A set of orthogonal versatile interacting peptide tags for imaging cellular proteins"

### Materials and Methods

*Chemicals*

All chemicals were acquired from commercial vendors: Millipore Sigma, Thermo Fisher Scientific, Fisher Scientific, Gold Bio, AK Scientific, and Lumiprobe. Chemicals were used as received without further purification.

*Generation of Plasmids*

The summary of all peptides and genetic constructs used in this work is presented in **Table S1**. Oligonucleotides were purchased from Integrated DNA Technologies (IDT); sequences are available upon request. Bacterial strains and vectors used in the present work are described in **Table S2**. New plasmids described in this work are available through Addgene.

The MiniR and MiniE expression plasmids were reported previously[1]. The pET28b plasmids for expression of TREER, TinyERRE, and PuRRRE were obtained using gene assembly PCR, as described[2]. Sequences were confirmed by Sanger sequencing (Genewiz). The pET28b(+)_PunyEEER plasmid was purchased (Genscript).

TOMM20-mCherry-N-10 (Addgene #55146) and mEmerald-Actin-C-18 (Addgene #53978) were obtained from Addgene. Vectors and the tag insert were amplified by PCR. Template DNA was removed by Dpn1 (NEB) digestion and PCR products were purified using Nucleospin Gel and PCR Clean-up kit (Takara). The insert and backbone were combined and ligated using Gibson Assembly Master Mix (NEB), following the manufacturer's instructions, and then transformed into NEB-5α cells (NEB). Transformed cells were selected on LB/kanamycin (50 μg/mL) plates and positive clones were confirmed by Sanger sequencing (Genewiz).

*Expression and Purification of Peptides*

Peptides were expressed from transformed *E. coli* BL21(DE3) cells (NEB) grown at 37 °C, 250 rpm in 2XYT media supplemented with kanamycin (50 μg/mL). The temperature was lowered to 30 °C before induction with 0.25 mM isopropyl β-D-1-thiogalactopyranoside (IPTG, GoldBio). After 4 hr, cells were harvested by centrifugation (4800xg, 4 °C) and pellets were stored at -20 °C prior to lysis.

Bacterial cells were lysed following an adapted protocol for the purification of recombinant protein from inclusion bodies[3]. The cell pellet was thawed on ice and resuspended in cell lysis buffer (100 mM NaCl, 1 mM EDTA, 50 mM Tris-Cl, pH 8.0 at 4 °C) supplemented with phenylmethylsulfonyl fluoride (PMSF, Thermo Fisher Scientific) and lysozyme (Thermo Fisher Scientific). After 20 min (4 °C rotation), deoxycholic acid (Fisher Scientific) was added before lysis by sonication (on ice). The lysate was clarified by centrifugation (15,000xg, 15 min, 4 °C) to obtain the soluble protein fraction. The pellet (with peptide in inclusion bodies) was washed twice with cell lysis buffer supplemented with 0.5% Triton X-100 (Millipore Sigma). The pellet was resuspended in Buffer B (6 M urea, 10 mM Tris-HCl, 100mM $NaH_2PO_4$, pH 8.0 at 4 °C) with PMSF (100 μM) and incubated (2 hr, room temperature [RT]). The urea fraction was clarified by centrifugation (15,000xg, 15 min, 4 °C) and then used for purification. Fractions were analyzed by SDS-PAGE; peptides were obtained from the urea-solubilized fraction.

Peptides were purified under denaturing conditions by immobilized metal affinity chromatography following The QIAexpressionist Handbook[4]. Peptides were bound to Ni-NTA resin in Buffer B supplemented with 10 mM imidazole (Thermo Fisher Scientific). The column was washed with 10 column volumes (CV) of Buffer B and 10 CV of Buffer C (6 M urea, 10 mM Tris-HCl, 100 mM $NaH_2PO_4$, pH 6.3 at 4 °C). The $His_6$-tagged peptides were eluted with 5 CV of Buffer E (6 M urea, 10 mM Tris-HCl, 100 mM $NaH_2PO_4$, pH 4.5 at 4 °C) and collected in 1 CV aliquots. The purification was analyzed by SDS-PAGE; purified peptides were in the Buffer E eluate. Peptides were concentrated (3 kDa MWCO centrifugal filter unit; Millipore Sigma) and buffer exchanged into 2 M TBS Urea (2 M urea, 10 mM Tris-HCl, 100 mM $NaH_2PO_4$, pH 8.0 at 4 °C) with 10 mM TCEP (Millipore Sigma). Purified peptide concentrations were determined by measuring the absorbance at 280 nm before storage at -20 °C in 2 M TBS Urea with 10% glycerol.

*SDS-PAGE analysis*

Peptide expression and purification was analyzed by SDS-PAGE on 12% Bis-Tris Criterion XT gels (Bio-Rad) in 1x MES buffer (Bio-Rad). Proteins were stained with Coomassie R-250 Brilliant Blue before imaging on a flatbed scanner (Epson Perfection Scanner V39).

*Circular Dichroism (CD) Spectroscopy*

Three replicates of each peptide were prepared by diluting to 5 µM in CD Buffer (12.5 mM $KH_2PO_4$ pH 7.4, 150 mM KCl, 1 mM EDTA)[5]. Peptides were incubated with 10 mM TCEP overnight at RT to reduce the peptides and increase solubility.

CD spectra were recorded using an AVIV 215 spectropolarimeter (AVIV Biomedical). CD spectra of individual peptides were measured at 5 µM total peptide concentration and mixtures of two peptides were measured at a 1:1 ratio (5 µM each; 10 µM total). Measurements were taken in millidegrees (mdeg) between 200-260 nm at 25 °C in 1 mm quartz cuvettes (Hellma 110-1-40) at 5 sec/nm averaging time with 1 nm steps. Molar ellipticity $[\theta]$ was calculated using Eq. 1. The mean residue molecular weight $(MWR)$ was found by the molecular weight of the peptides divided by the number of amino acid residues. Cell path length $(l)$ was 0.1 cm and $c$ is the total peptide concentration in g/L.

$$[\theta] = \frac{mdeg \times MWR}{10 \times l \times c} \qquad \text{Eq. 1}$$

Denaturation of the heterodimers was measured in guanidine hydrochloride (GdnHCl, Thermo Fisher Scientific) in CD Buffer supplemented with 10 mM TCEP. Peptides were prepared in 0 to 6 M GdnHCl at 0.5 M increments. Samples were analyzed with 30 sec reads at 222 nm. The GdnHCl denaturation curves were analyzed following published methods[6]. The mean residue molar ellipticity $[\theta]$ was calculated (Eq. 2), where the observed ellipticity $(\theta_{obs})$ was found by taking the y-intercept of the linear regression of the denaturation gradient. The y-intercept of the buffer blank was subtracted from the y-intercept of the peptide samples.

$$[\theta] = \frac{\theta_{obs} \times MWR}{10 \times l \times c} \qquad \text{Eq. 2}$$

A two-state model was used to calculate the fraction folded $\left(F_f\right)$ (Eq. 3), where $[\theta]_f$ is the ellipticity of the folded state and $[\theta]_d$ is the ellipticity of the denatured state. The 100% folded state of each peptide pair was observed between 0 and 1.0 M GdnHCl. As a result, the $[\theta]$ of all replicates from 0-1.0 M were averaged to determine the $[\theta]_f$. An average of all three replicates at 6 M was assumed to represent a 100% denatured state. The unfolded fraction $(F_u)$ was calculated using Eq. 4.

$$F_f = \frac{[\theta]-[\theta]_d}{[\theta]_f-[\theta]_d} \qquad \text{Eq. 3}$$

$$F_u = 1 - F_f \qquad \text{Eq. 4}$$

The free energy of unfolding, $\Delta G_D$, was calculated using Eq. 5, where $R$ is the molar gas constant (8.314 J/mol*K), $T$ is the temperature in kelvin (298 K), and $P_t$ is the total peptide concentration (mol).

$$\Delta G_D = -RT \ln\left(\frac{2P_t(F_u)^2}{F_f}\right) \qquad \text{Eq. 5}$$

Linear extrapolation was used to calculate the free energy of unfolding without GdnHCl ($\Delta G^{H_2O}$) using Eq. 6 and assuming a linear relationship between the free energy of unfolding and concentration of denaturant [GdnHCl]. The $K_{D(app)}$ was calculated using Eq. 7.

$$\Delta G_D = \Delta G^{H_2O} - m[GdnHCl] \qquad \text{Eq. 6}$$

$$\Delta G^{H_2O} = -RT ln K_{D(app)} \qquad \text{Eq. 7}$$

*Generation of Probe Peptides by Thiol-Maleimide Chemistry*

The probe peptides used in this work (**Table S4**) were labeled using thiol-maleimide chemistry with azide quenching of TCEP[7]. Reactions containing at least 100 μM of peptide were mixed with 250 molar equivalents TCEP in Labeling Buffer (0.5 M Tris-HCl, 150 mM NaCl, pH 7.2 at 32 °C). After 45 min at RT, 1250 equivalents of 1,14-diazido-3,6,9,12-tetraoxatetradecane (CAS 182760-73-2, AK Scientific) was added to quench excess TCEP (1h at 37 °C with agitation). Next, maleimide dye (20 equivalents, **Table S3**) was added and the mixture was incubated overnight at 37 °C with agitation. Free fluorophore was removed through purification by immobilized metal affinity chromatography (Ni-NTA agarose; Qiagen). The labeling reaction was bound to resin in 2M Urea-T (20 mM Tris-HCl, 150 mM NaCl, 2 M urea pH 7.4) for 1 hr (4 °C). The column was washed with 60 CV 2M Urea-T and eluted in 2M Urea-T with 250 mM imidazole (2 CV) and 500 mM imidazole (3 CV). Fractions with fluorophore-labeled peptide were combined and the concentration and degree of labeling (% labeled) were determined using Thermo Fisher's protocol: TR0031-Calc-FP-ratios.pdf (thermofisher.com). Probe peptides were stored in 2M Urea-T with 10% glycerol at -20 °C.

*Mammalian Cell Culture*

Chinese hamster ovary (CHO) TRVb (ΔTfR1 ΔTfR2) cells were provided by Prof. Timothy E. McGraw (Cornell University, New York). These cells do not express transferrin receptor 1 (TfR1) or transferrin receptor 2 (TfR2)[8]. U-2 OS cells were purchased from ATCC (Cat. # HTB-96). CHO TRVb and U-2 OS cells were maintained in Ham F-12 media (Gibco) with 5% fetal bovine serum (FBS, Gibco) or McCoy's 5A media (Gibco) with 10% FBS, respectively. Cells were grown in a humidified incubator at 37 °C with 5% $CO_2$. Cells were passaged (80-90% confluency) using 0.25% trypsin, 1 mM EDTA (Gibco). Cells were plated in 10 $cm^2$ polystyrene dishes at $4x10^5$ cells/dish (CHO) or $5x10^5$ cells/dish (U-2 OS). Cultured cells were assessed routinely for mycoplasma contamination through PCR and Hoechst staining.

*Cell Transfection*

All labeling experiments used transiently transfected cells. Cells were seeded ($3x10^4$ cells/well) into 8-well, #1.5 chambered coverslips (Ibidi, Cat. # 80806) and grown to 70-90% confluency. Cells were transfected with TransIT-2020 (Mirus Bio) according to the manufacturer's instructions. All transfections were performed using a total DNA-to-reagent ratio of 2:1. For single-target imaging (TfR1-tagged constructs), CHO TRVb cells were transfected with 180 ng of vector DNA. For two-target imaging, cells (CHO TRVb or U-2 OS) were transfected with vector DNA as follows: 180 ng pcDNA3.1_TfR1, pcDNA3.1_TfR1-PunyEEER; 60 ng mEmerald-H2B-6, H2B-6-MiniR-mEm; 120 ng mEmeral-Actin-C-18, mEm-TREER-Actin; and/or 60 ng mCherry-TOMM20-N-10, TOMM20-PuRRRE. For three-target imaging, transfections were done with 60 ng mCherry-TOMM20-N-10, TOMM20-PuRRRE; 60 ng mEmerald-H2B-6, H2B-6-MiniR; and 60 ng mEmeral-Actin-C-18, TREER-Actin. Cells were labeled ~24 hr after transfection. All labeling experiments were repeated a minimum of three times.

*Labeling of VIP-Tagged TfR1 in Live Cells for Florescence Microscopy (FM)*

After transfection, CHO TRVb cells were blocked (30 min, 37 °C) with live-cell block buffer (LCB: Ham F12 medium, 5% FBS, 6% bovine serum albumin [BSA]) and then chilled on ice. Cells were incubated (30 min, 4 °C) with 100 nM Cy5-labeled probe peptide and 50 ng/mL transferrin-AF488 (Tf-488, Thermo Fisher Scientific) in cold LCB. Cells were then either further incubated for 10 min at 37 °C or fixed immediately. For fixation, cells were washed twice with ice-cold phosphate buffered saline (PBS) and incubated (10 min, 4 °C) in 4% paraformaldehyde (PFA) with 5 µg/mL Hoechst 33342 nuclear stain (Thermo Fisher Scientific). Cells were washed thrice with PBS, overlaid with mounting media (80% glycerol, 20 mM Tris, 0.01% sodium azide), and imaged.

*Labeling of Intracellular VIP-Tagged Proteins in Fixed Cells for FM*

Transfected cells were fixed with 4% PFA (10 min, RT) and permeabilized (0.1% Triton-X 100) for 15 min. Fixation of cells expressing TOMM20 was done at 37 °C to preserve mitochondrial morphology. Cells were blocked (5% BSA in PBS) for 30 min. For two-color and three-color labeling, cells were treated simultaneously with probe peptides (100 nM each) and either 0.5 µg/mL 4′,6-diamidino-2-phenylindole (DAPI, Fisher Scientific) or 5 µg/mL Hoechst 33342 in 5% BSA for 30 min at RT. Cells were washed, overlaid with mounting media, and imaged.

*Fluorescence Microscopy and Image Analysis*

All micrographs were acquired on a spinning disk confocal microscope (Yokogawa CSU-X1 on a Zeiss Axio Observer) using a 63X Plan-Apochromat oil-immersion objective lens (N.A. 1.4, Zeiss). All samples within an experiment were imaged using identical acquisition settings (laser intensity, exposure time, filters, etc.) and a minimum of three fields of view were collected per sample. The following excitation (ex) lasers and emission (em) filters were used for the specified fluorophores: DAPI/Hoechst 33342: ex 405 nm, em 405/50 nm; mEm, AF488: ex 488 nm, em 525/50 nm; Cy3, mCherry: ex 561 nm, em 629/62 nm; Cy5: ex 638 nm, em 690/50 nm.

Micrographs were processed using Fiji[9]. Brightness and contrast levels for each channel were set identically across all images within an experiment. Merged micrographs were generated in Fiji by overlaying each channel within an image. Micrographs were cropped to provide enlarged views of representative cells. Fluorescence colocalization was analyzed in Zen software (Zeiss) using manual thresholding based on single color controls and a region of interest defined around the representative cell (**Figure S3**). Colocalization was assessed through the Pearson's correlation coefficient (PCC) statistic[10]. Figures were prepared in Inkscape (https://inkscape.org/).

### Supplementary Figures.

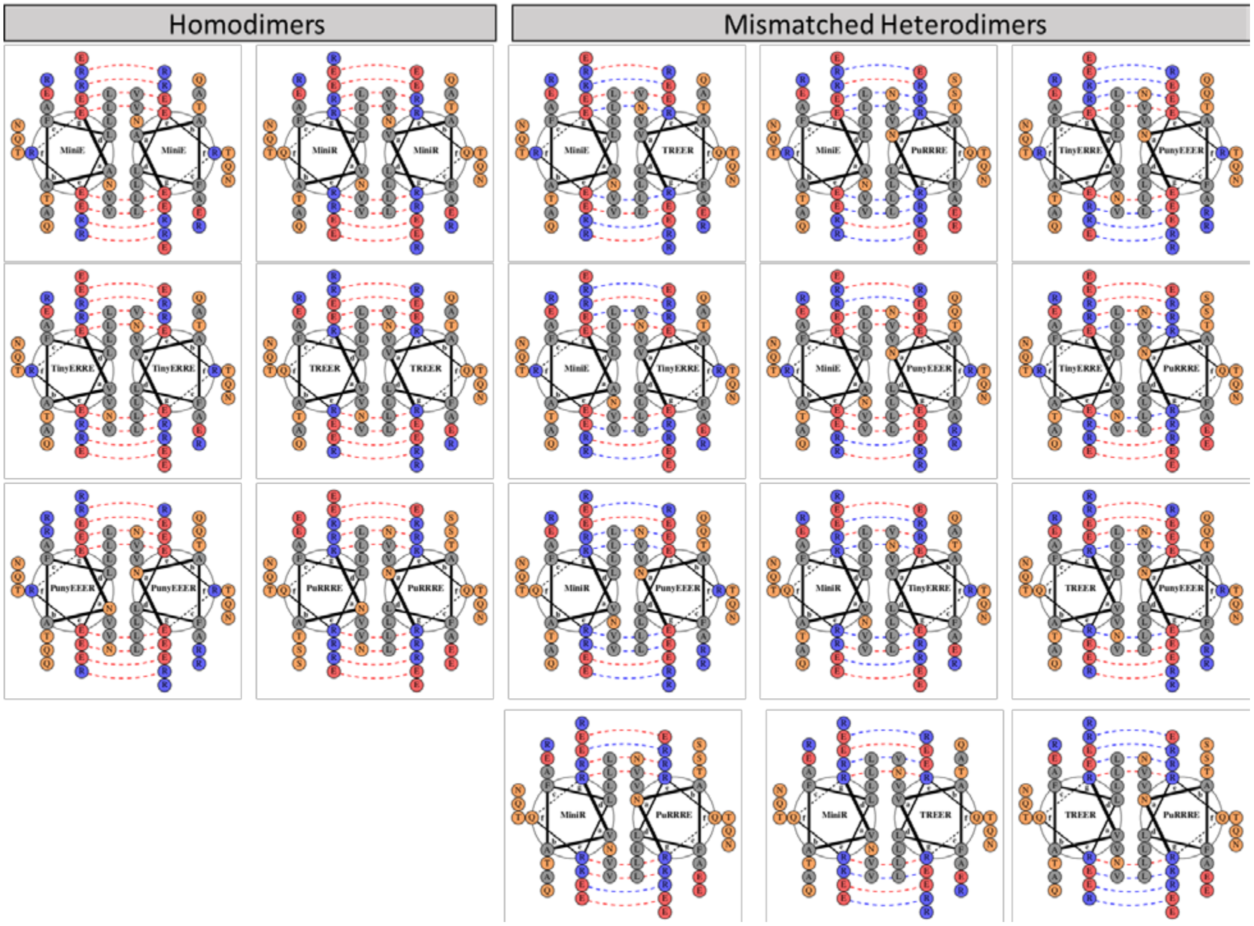


**Figure S1. Helical wheel diagrams of disfavored homodimeric coiled coils and mismatched heterodimeric coiled coils.** The three VIP tags (MiniVIPER, TinyVIPER, and PunyVIPER) are heterodimers that all contain Asn-Asn matches and 8 favorable salt bridges (*e* and *g* positions), see **Fig. 2**. In contrast, all six homodimers include 8 unfavorable (repulsive) interactions at the *e* and *g* positions. Only parallel dimers are shown because anti-parallel dimers are disfavored due to the lack of Asn-Asn matching at the *a* position. The heterodimeric (mismatched) coiled coils depicted here have a mixture of favorable (E—R) salt bridges (blue dashed lines) and repellent (E—E or R—R) interactions (red dashed lines). Amino acids are color-coded (red = negative charge; blue = positive charge; gray = hydrophobic; yellow = hydrophilic). Helical wheel diagrams were generated using DrawCoil1.0 (https://grigoryanlab.org/drawcoil/).

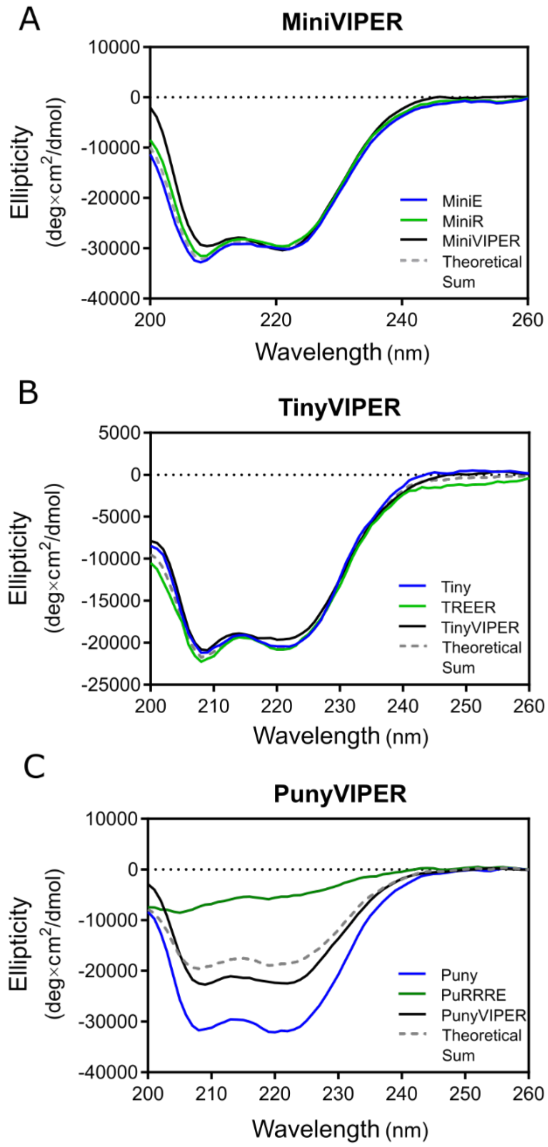


**Figure S2. Secondary structure of MiniVIPER, TinyVIPER, and PunyVIPER.** CD spectra between 200-260 nm of peptide monomers (blue and green), coiled-coil dimers in a 1:1 molar ratio of each monomer (black), and the theoretical sum of the monomers (gray dashed). Spectra were measured for MiniVIPER (A), TinyVIPER (B), and PunyVIPER (C).

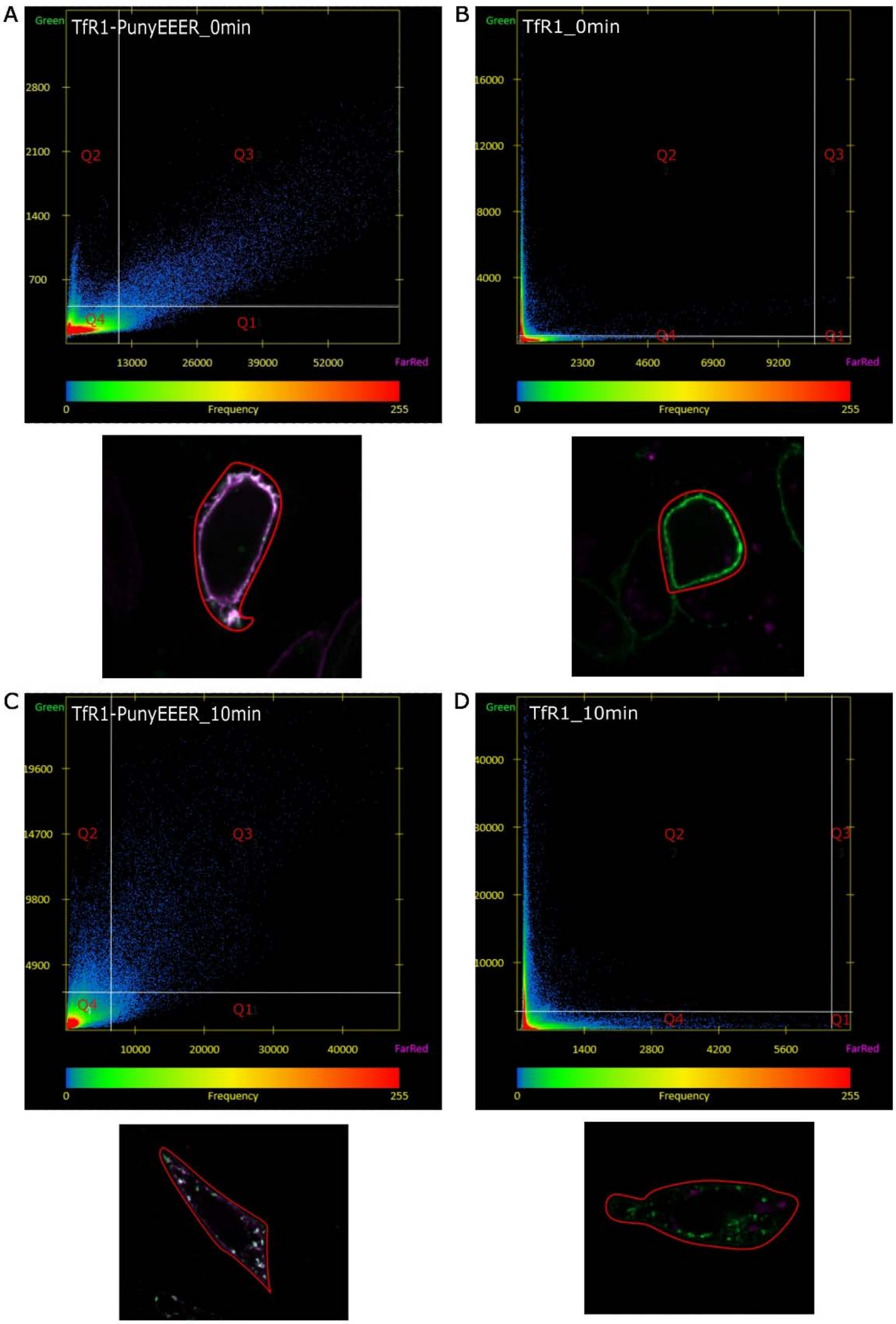


**Figure S3. Regions of interest and channel intensity scatter plots for colocalization analysis.** Regions of interest (red outline) in images of CHO TRVb cells expressing TfR1-PunyEEER (A, C) or untagged TfR1 (B, D) were defined for the analysis of colocalization between Tf-488 (Green) and PuRRRE-Cy5 (FarRed) signals. Channel intensity scatter plots display pixels defined as FarRed only (Q1), Green only (Q2), both Green and FarRed (colocalized, Q3), or background (Q4). Pearson's correlation coefficients estimating the linearity of these distributions were calculated and reported in **Fig. 4**. Equivalent analyses were completed for all images in **Fig. 4**.

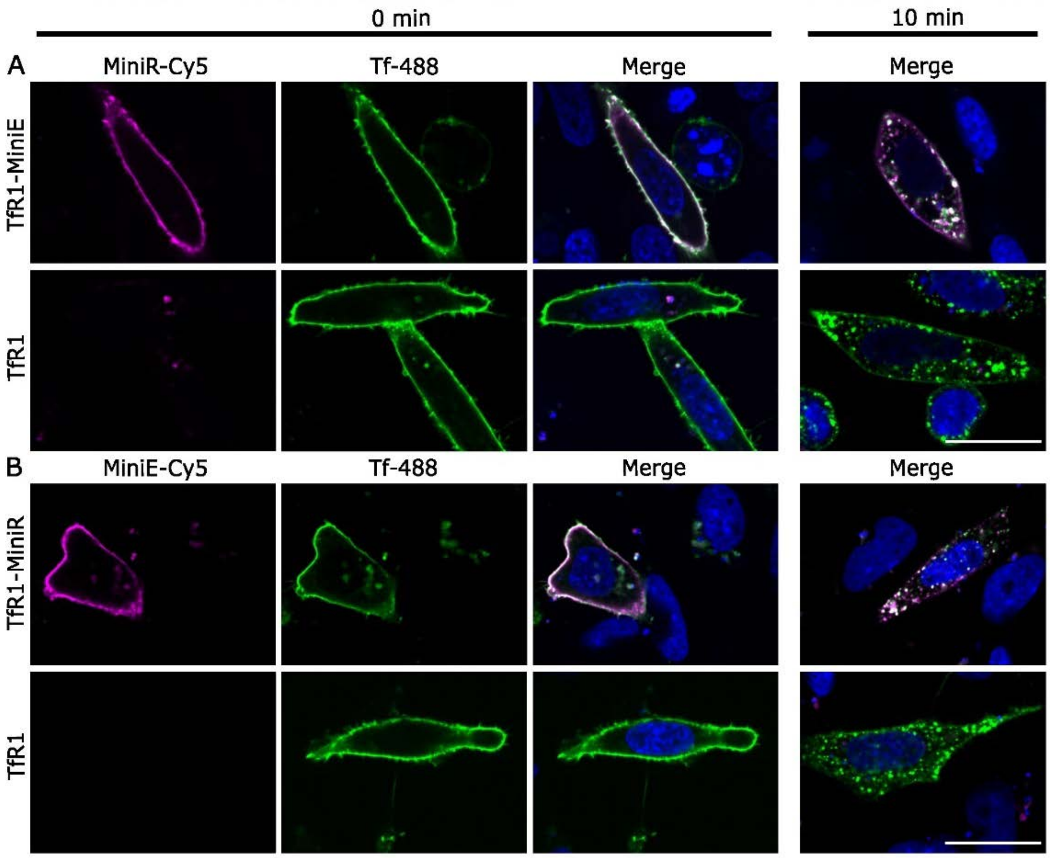


**Figure S4. MiniVIPER-tagged TfR1 receptor localizes to the membrane, binds Tf, and trafficks by endocytosis.** CHO TRVb cells expressing MiniVIPER-tagged TfR1 were labeled live (4 °C) with 100 nM Cy5-conjugated probe peptide and 50 ng/mL Tf-488. Cells were fixed immediately after labeling (0 min) or after incubation at 37 °C to allow receptor-ligand internalization (10 min). (A) Cells expressing TfR1-MiniE or untagged TfR1 labeled with MiniR-Cy5 and Tf-488. (B) Cells expressing TfR1-MiniR or TfR1 labeled with MiniE-Cy5 and Tf-488. Micrographs are false-colored (Cy5, magenta; Tf-488, green; Hoechst, blue) and green-magenta overlap appears white in the merge. Scale bars represent 20 μm.

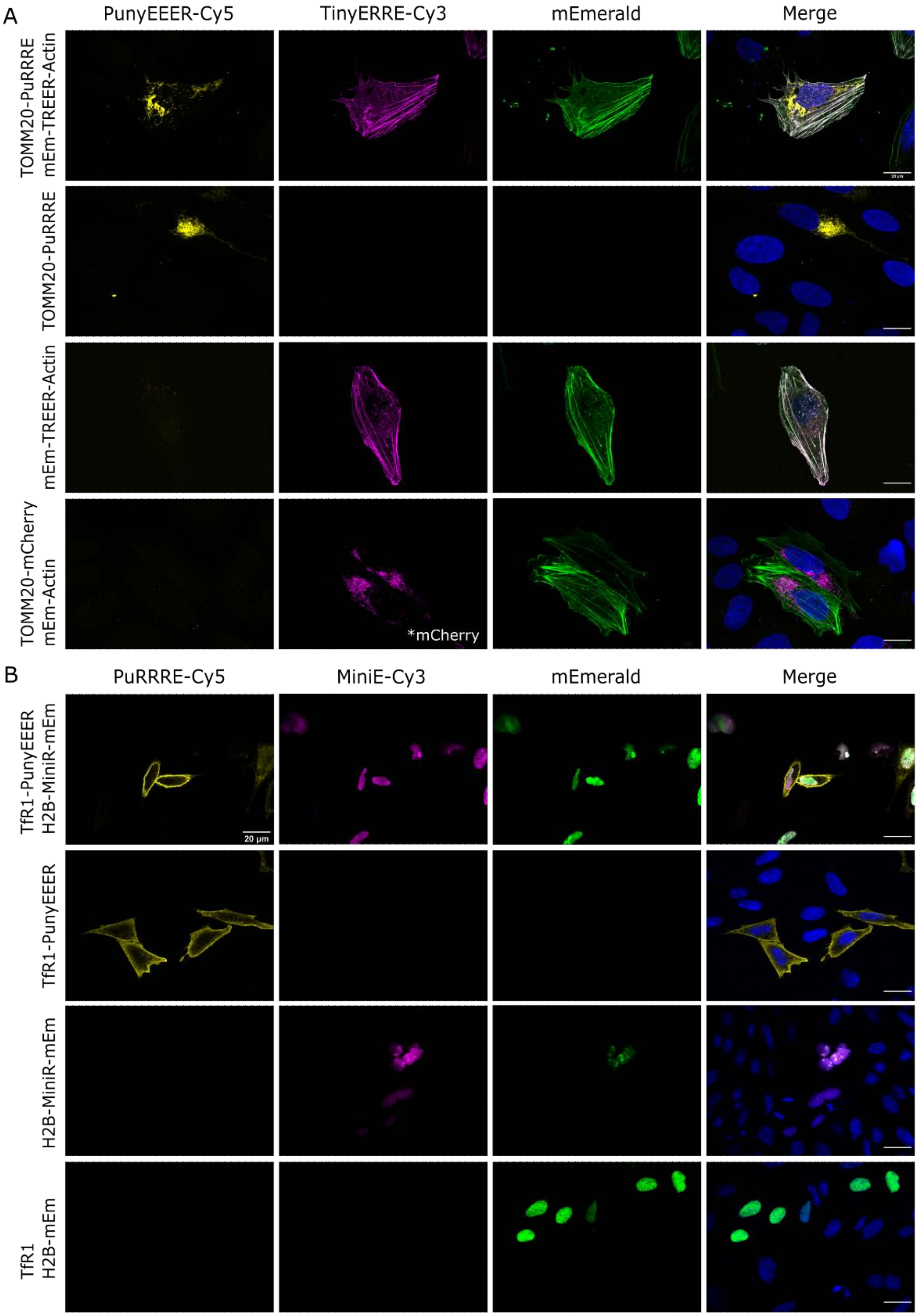


**Figure S5. VIP tags enable specific labeling of two protein targets in cells.** (A) U-2 OS cells expressing mEm-TREER-Actin, TOMM20-PuRRRE, TOMM20-mCherry, and/or mEm-Actin were simultaneously treated with 100 nM PunyEEER-Cy5 and 100 nM TinyERRE-Cy3. Cells were imaged to detect fluorescently-labeled mitochondria and actin filaments. Micrographs are presented as a maximum intensity projection of a confocal z-stack. (B) CHO TRVb cells expressing TfR1-PunyEEER, H2B-MiniR-mEm, TfR1, and/or H2B-mEm were treated with 100 nM PuRRRE-Cy5 (live) and 100 nM MiniE-Cy3 (post-fixation). Micrographs are false-colored (Cy3/mCherry, magenta; Cy5, yellow; mEm, green) and the merge includes DAPI/Hoechst nuclear stain (blue). Scale bars represent 20 µm.

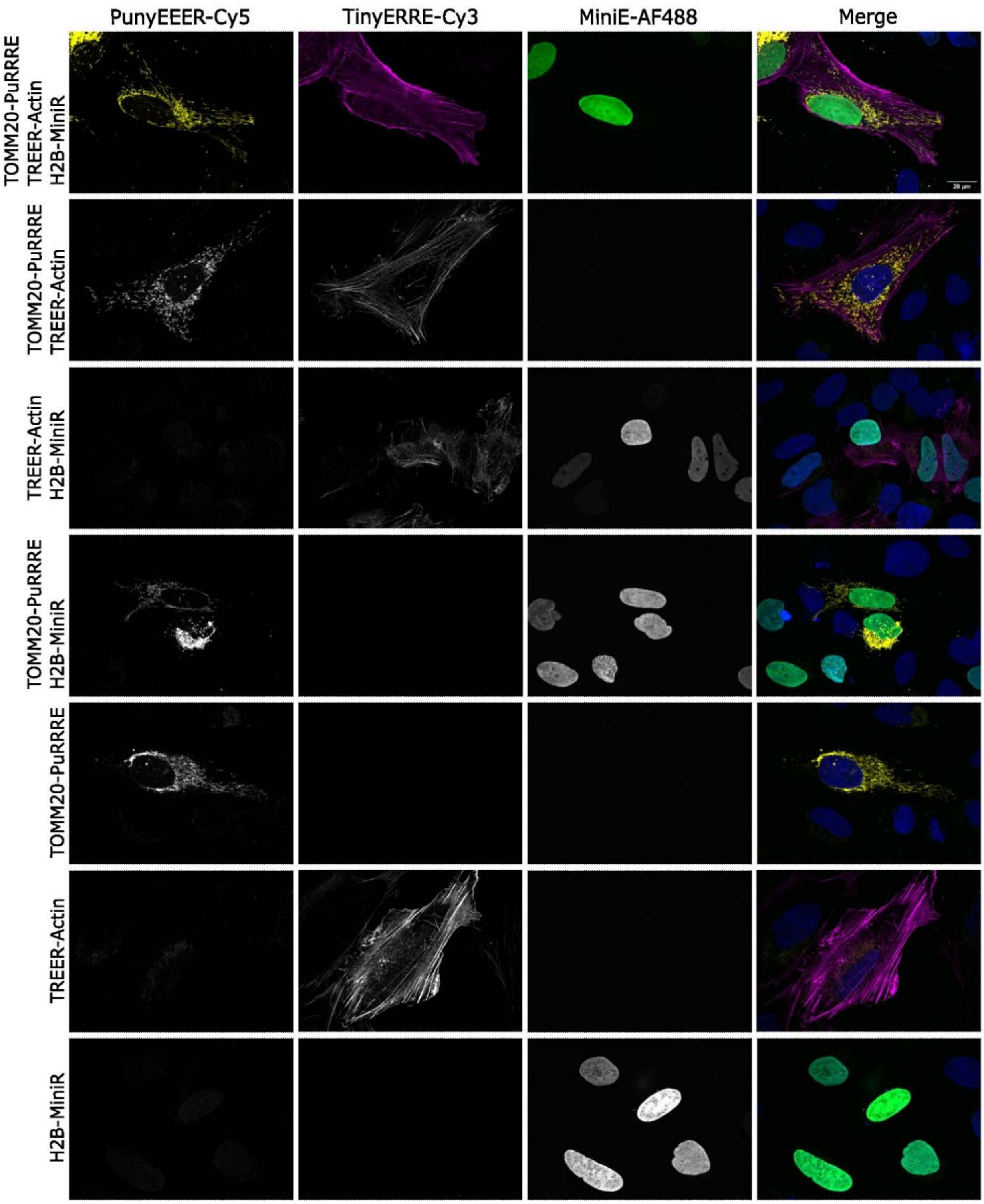


**Figure S6**. **MiniVIPER, TinyVIPER, and PunyVIPER enable simultaneous imaging of three distinct protein targets in cells.** U-2 OS cells expressing TREER-Actin, H2B-MiniR and/or TOMM20-PuRRRE were fixed and then treated simultaneously with 100 nM each of PunyEEER-Cy5, TinyERRE-Cy3, and MiniE-AF488. Cells were imaged by confocal FM imaging and micrographs represent maximum intensity projections from z-stacks. Micrographs are false-colored (Cy5, yellow; Cy3, magenta; AF488, green; DAPI, blue). Scale bars represent 20 µm.

### Supplementary Tables.

#### Table S1: Summary of genetic constructs.

| Peptide / Protein | Amino acid sequence<br>Key: **Coil tag**; DBCO tag; linker; mEmerald; mCherry | MW (Da) | Vector Name |
|---|---|---|---|
| MiniE | MGGS**LEIEAAFLERENTALETRVAELRQRVQRLRNEYGPL**GGGAAAWGLCYPW<br>VYGLEHHHHHH | 7322.2 | pET28(+)_MiniE<br>Addgene: 181985 |
| MiniR | MGGS**LEIRVAFLRQRNTALRTEVAELEQEVQRLENRYGPL**GGGAAAWGLCYPWVYGLEHH<br>HHHH | 7349.3 | pET28(+)_MiniR<br>Addgene: 181984 |
| TinyERRE | MGGS**LEIEVAFLERRVTALRTRNAELRQEVQRLENEYGPL**GGGAAAWGLCYPWVYGLEHHH<br>HHH | 7350.2 | pET28(+)_TinyERRE |
| TREER | MGGS**LEIRVAFLRQEVTALETENAELEQRVQRLRNRYGPL**GGGAAAWGLCYPWVYGLEHH<br>HHHH | 7349.3 | pET28(+)_TREER |
| PunyEEER | MGGS**LEIENAFLEREVTALETEVQRLEQRNQRLRNRYGPL**GSWGLCYPWVYGLEHHHHHH | 7182.0 | pET28(+)_PunyEEER |
| PuRRRE | MGGS**LEIRNAFLRQRVTALRTRVSELRQENSELENEYGPL**GSWGLCYPWVYGLEHHHHHH | 7098.9 | pET28(+)_PuRRRE |
| Transferrin Receptor 1 (TfR1) | MMDQARSAFSNLFGGEPLSYTRFSLARQVDGDNSHVEMKLAADEEENADNNMKASVRKPKR<br>FNGRLCFAAIALVIFFLIGFMSGYLGYCKRVEQKEECVKLAETEETDKSETMETEDVPTSSRLY<br>WADLKTLLSEKLNSIEFADTIKQLSQNTYTPREAGSQKDESLAYYIENQFHEFKFSKVWRDEHY<br>VKIQVKSSIGQNMVTIVQSNGNLDPVESPEGYVAFSKPTEVSGKLVHANFGTKKDFEELSYSV<br>NGSLVIVRAGEITFAEKVANAQSFNAIGVLIYMDKNKFPVVEADLALFGHAHLGTGDPYTPGFP<br>SFNHTQFPPSQSSGLPNIPVQTISRAAAEKLFGKMEGSCPARWNIDSSCKLELSQNQNVKLIVK<br>NVLKERRILNIFGVIKGYEEPDRYVVVGAQRDALGAGVAAKSSVGTGLLLKLAQVFSDMISKDG<br>FRPSRSIIFASWTAGDFGAVGATEWLEGYLSSLHLKAFTYINLDKVVLGTSNFKVSASPLLYTLM<br>GKIMQDVKHPVDGKSLYRDSNWISKVEKLSFDNAAYPFLAYSGIPAVSFCFCEDADYPYLGTR<br>LDTYEALTQKVPQLNQMVRTAAEVAGQLIIKLTHDVELNLDYEMYNSKLLSFMKDLNQFKTDIR<br>DMGLSLQWLYSARGDYFRATSRLTTDFHNAEKTNRFVMREINDRIMKVEYHFLSPYVSPRESP<br>FRHIFWGSGSHTLSALVENLKLRQKNITAFNETLFRNQLALATWTIQGVANALSGDIWNIDNEF* | 85731.4 | pcDNA3.1_TfR1 |
| TfR1-MiniE | MMDQARSAFSNLFGGEPLSYTRFSLARQVDGDNSHVEMKLAADEEENADNNMKASVRKPKR<br>FNGRLCFAAIALVIFFLIGFMSGYLGYCKRVEQKEECVKLAETEETDKSETMETEDVPTSSRLY<br>WADLKTLLSEKLNSIEFADTIKQLSQNTYTPREAGSQKDESLAYYIENQFHEFKFSKVWRDEHY<br>VKIQVKSSIGQNMVTIVQSNGNLDPVESPEGYVAFSKPTEVSGKLVHANFGTKKDFEELSYSV<br>NGSLVIVRAGEITFAEKVANAQSFNAIGVLIYMDKNKFPVVEADLALFGHAHLGTGDPYTPGFP<br>SFNHTQFPPSQSSGLPNIPVQTISRAAAEKLFGKMEGSCPARWNIDSSCKLELSQNQNVKLIVK<br>NVLKERRILNIFGVIKGYEEPDRYVVVGAQRDALGAGVAAKSSVGTGLLLKLAQVFSDMISKDG<br>FRPSRSIIFASWTAGDFGAVGATEWLEGYLSSLHLKAFTYINLDKVVLGTSNFKVSASPLLYTLM<br>GKIMQDVKHPVDGKSLYRDSNWISKVEKLSFDNAAYPFLAYSGIPAVSFCFCEDADYPYLGTR<br>LDTYEALTQKVPQLNQMVRTAAEVAGQLIIKLTHDVELNLDYEMYNSKLLSFMKDLNQFKTDIR<br>DMGLSLQWLYSARGDYFRATSRLTTDFHNAEKTNRFVMREINDRIMKVEYHFLSPYVSPRESP<br>FRHIFWGSGSHTLSALVENLKLRQKNITAFNETLFRNQLALATWTIQGVANALSGDIWNIDNEF<br>GSGSGSTGM**LEIEAAFLERENTALETRVAELRQRVQRLRNEYGPL**GGGTG* | 91049.3 | pcDNA3.1_TfR1-MiniE |
| TfR1-MiniR | MMDQARSAFSNLFGGEPLSYTRFSLARQVDGDNSHVEMKLAADEEENADNNMKASVRKPKR<br>FNGRLCFAAIALVIFFLIGFMSGYLGYCKRVEQKEECVKLAETEETDKSETMETEDVPTSSRLY<br>WADLKTLLSEKLNSIEFADTIKQLSQNTYTPREAGSQKDESLAYYIENQFHEFKFSKVWRDEHY<br>VKIQVKSSIGQNMVTIVQSNGNLDPVESPEGYVAFSKPTEVSGKLVHANFGTKKDFEELSYSV<br>NGSLVIVRAGEITFAEKVANAQSFNAIGVLIYMDKNKFPVVEADLALFGHAHLGTGDPYTPGFP<br>SFNHTQFPPSQSSGLPNIPVQTISRAAAEKLFGKMEGSCPARWNIDSSCKLELSQNQNVKLIVK<br>NVLKERRILNIFGVIKGYEEPDRYVVVGAQRDALGAGVAAKSSVGTGLLLKLAQVFSDMISKDG<br>FRPSRSIIFASWTAGDFGAVGATEWLEGYLSSLHLKAFTYINLDKVVLGTSNFKVSASPLLYTLM<br>GKIMQDVKHPVDGKSLYRDSNWISKVEKLSFDNAAYPFLAYSGIPAVSFCFCEDADYPYLGTR<br>LDTYEALTQKVPQLNQMVRTAAEVAGQLIIKLTHDVELNLDYEMYNSKLLSFMKDLNQFKTDIR<br>DMGLSLQWLYSARGDYFRATSRLTTDFHNAEKTNRFVMREINDRIMKVEYHFLSPYVSPRESP<br>FRHIFWGSGSHTLSALVENLKLRQKNITAFNETLFRNQLALATWTIQGVANALSGDIWNIDNEF<br>GSGSGSTGMGSS**LEIRVAFLRQRNTALRTEVAELEQEVQRLENRYGPL**TG* | 91136.4 | pcDNA3.1_TfR1-MiniR<br>Addgene: 181986 |
| TfR1-TREER | MMDQARSAFSNLFGGEPLSYTRFSLARQVDGDNSHVEMKLAADEEENADNNMKASVRKPKR<br>FNGRLCFAAIALVIFFLIGFMSGYLGYCKRVEQKEECVKLAETEETDKSETMETEDVPTSSRLY<br>WADLKTLLSEKLNSIEFADTIKQLSQNTYTPREAGSQKDESLAYYIENQFHEFKFSKVWRDEHY<br>VKIQVKSSIGQNMVTIVQSNGNLDPVESPEGYVAFSKPTEVSGKLVHANFGTKKDFEELSYSV<br>NGSLVIVRAGEITFAEKVANAQSFNAIGVLIYMDKNKFPVVEADLALFGHAHLGTGDPYTPGFP<br>SFNHTQFPPSQSSGLPNIPVQTISRAAAEKLFGKMEGSCPARWNIDSSCKLELSQNQNVKLIVK<br>NVLKERRILNIFGVIKGYEEPDRYVVVGAQRDALGAGVAAKSSVGTGLLLKLAQVFSDMISKDG<br>FRPSRSIIFASWTAGDFGAVGATEWLEGYLSSLHLKAFTYINLDKVVLGTSNFKVSASPLLYTLM<br>GKIMQDVKHPVDGKSLYRDSNWISKVEKLSFDNAAYPFLAYSGIPAVSFCFCEDADYPYLGTR<br>LDTYEALTQKVPQLNQMVRTAAEVAGQLIIKLTHDVELNLDYEMYNSKLLSFMKDLNQFKTDIR<br>DMGLSLQWLYSARGDYFRATSRLTTDFHNAEKTNRFVMREINDRIMKVEYHFLSPYVSPRESP<br>FRHIFWGSGSHTLSALVENLKLRQKNITAFNETLFRNQLALATWTIQGVANALSGDIWNIDNEF<br>GSGSGSTG**LEIRVAFLRQEVTALETENAELEQRVQRLRNRYGPL**TG* | 90774.0 | pcDNA3.1_TfR1-TREER |
| TfR1-TinyERRE | MMDQARSAFSNLFGGEPLSYTRFSLARQVDGDNSHVEMKLAADEEENADNNMKASVRKPKR<br>FNGRLCFAAIALVIFFLIGFMSGYLGYCKRVEQKEECVKLAETEETDKSETMETEDVPTSSRLY<br>WADLKTLLSEKLNSIEFADTIKQLSQNTYTPREAGSQKDESLAYYIENQFHEFKFSKVWRDEHY<br>VKIQVKSSIGQNMVTIVQSNGNLDPVESPEGYVAFSKPTEVSGKLVHANFGTKKDFEELSYSV<br>NGSLVIVRAGEITFAEKVANAQSFNAIGVLIYMDKNKFPVVEADLALFGHAHLGTGDPYTPGFP<br>SFNHTQFPPSQSSGLPNIPVQTISRAAAEKLFGKMEGSCPARWNIDSSCKLELSQNQNVKLIVK<br>NVLKERRILNIFGVIKGYEEPDRYVVVGAQRDALGAGVAAKSSVGTGLLLKLAQVFSDMISKDG<br>FRPSRSIIFASWTAGDFGAVGATEWLEGYLSSLHLKAFTYINLDKVVLGTSNFKVSASPLLYTLM<br>GKIMQDVKHPVDGKSLYRDSNWISKVEKLSFDNAAYPFLAYSGIPAVSFCFCEDADYPYLGTR<br>LDTYEALTQKVPQLNQMVRTAAEVAGQLIIKLTHDVELNLDYEMYNSKLLSFMKDLNQFKTDIR<br>DMGLSLQWLYSARGDYFRATSRLTTDFHNAEKTNRFVMREINDRIMKVEYHFLSPYVSPRESP<br>FRHIFWGSGSHTLSALVENLKLRQKNITAFNETLFRNQLALATWTIQGVANALSGDIWNIDNEF<br>GSGSGSTG**LEIEVAFLERRVTALRTRNAELRQEVQRLENEYGPL**TG* | 90775.0 | pcDNA3.1_TfR1-TinyERRE |
| TfR1- | MMDQARSAFSNLFGGEPLSYTRFSLARQVDGDNSHVEMKLAADEEENADNNMKASVRKPKR | 90847.0 | pcDNA3.1_TfR1- |

| | | | |
|---|---|---|---|
| PunyEEER | FNGRLCFAAIALVIFFLIGFMSGYLGYCKRVEQKEECVKLAETEETDKSETMETEDVPTSSRLY<br>WADLKTLLSEKLNSIEFADTIKQLSQNTYTPREAGSQKDESLAYYIENQFHEFKFSKVWRDEHY<br>VKIQVKSSIGQNMVTIVQSNGNLDPVESPEGYVAFSKPTEVSGKLVHANFGTKKDFEELSYSV<br>NGSLVIVRAGEITFAEKVANAQSFNAIGVLIYMDKNKFPVVEADLALFGHAHLGTGDPYTPGFP<br>SFNHTQFPPSQSSGLPNIPVQTISRAAAEKLFGKMEGSCPARWNIDSSCKLELSQNQNVKLIVK<br>NVLKERRILNIFGVIKGYEEPDRYVVVGAQRDALGAGVAAKSSVGTGLLLKLAQVFSDMISKDG<br>FRPSRSIIFASWTAGDFGAVGATEWLEGYLSSLHLKAFTYINLDKVVLGTSNFKVSASPLLYTLM<br>GKIMQDVKHPVDGKSLYRDSNWISKVEKLSFDNAAYPFLAYSGIPAVSFCFCEDADYPYLGTR<br>LDTYEALTQKVPQLNQMVRTAAEVAGQLIIKLTHDVELNLDYEMYNSKLLSFMKDLNQFKTDIR<br>DMGLSLQWLYSARGDYFRATSRLTTDFHNAEKTNRFVMREINDRIMKVEYHFLSPYVSPRESP<br>FRHIFWGSGSHTLSALVENLKLRQKNITAFNETLFRNQLALATWTIQGVANALSGDIWNIDNEF<br>GSGSGSTG**LEIENAFLEREVTALETEVQRLEQRNQRLRNRYGPL**TG* | | PunyEEER |
| TfR1-PuRRRE | MMDQARSAFSNLFGGEPLSYTRFSLARQVDGDNSHVEMKLAADEEENADNNMKASVRKPKR<br>FNGRLCFAAIALVIFFLIGFMSGYLGYCKRVEQKEECVKLAETEETDKSETMETEDVPTSSRLY<br>WADLKTLLSEKLNSIEFADTIKQLSQNTYTPREAGSQKDESLAYYIENQFHEFKFSKVWRDEHY<br>VKIQVKSSIGQNMVTIVQSNGNLDPVESPEGYVAFSKPTEVSGKLVHANFGTKKDFEELSYSV<br>NGSLVIVRAGEITFAEKVANAQSFNAIGVLIYMDKNKFPVVEADLALFGHAHLGTGDPYTPGFP<br>SFNHTQFPPSQSSGLPNIPVQTISRAAAEKLFGKMEGSCPARWNIDSSCKLELSQNQNVKLIVK<br>NVLKERRILNIFGVIKGYEEPDRYVVVGAQRDALGAGVAAKSSVGTGLLLKLAQVFSDMISKDG<br>FRPSRSIIFASWTAGDFGAVGATEWLEGYLSSLHLKAFTYINLDKVVLGTSNFKVSASPLLYTLM<br>GKIMQDVKHPVDGKSLYRDSNWISKVEKLSFDNAAYPFLAYSGIPAVSFCFCEDADYPYLGTR<br>LDTYEALTQKVPQLNQMVRTAAEVAGQLIIKLTHDVELNLDYEMYNSKLLSFMKDLNQFKTDIR<br>DMGLSLQWLYSARGDYFRATSRLTTDFHNAEKTNRFVMREINDRIMKVEYHFLSPYVSPRESP<br>FRHIFWGSGSHTLSALVENLKLRQKNITAFNETLFRNQLALATWTIQGVANALSGDIWNIDNEF<br>GSGSGSTG**LEIRNAFLRQRVTALRTRVSELRQENSELENEYGPL*** | 90605.8 | pcDNA3.1_TfR1-PuRRRE |
| H2B-mEmerald | MPEPAKSAPAPKKGSKKAVTKAQKKGGKKRKRSRKESYSIYVYKVLKQVHPDTGISSKAMGIM<br>NSFVNDIFERIAGEASRLAHYNKRSTITSREIQTAVRLLLPGELAKHAVSEGTKAITKYTSAKDPP<br>VATMVSKGEELFTGVVPILVELDGDVNGHKFSVSGEGEGDATYGKLTLKFICTTGKLPVPWPT<br>LVTTLTYGVQCFARYPDHMKQHDFFKSAMPEGYVQERTIFFKDDGNYKTRAEVKFEGDTLVN<br>RIELKGIDFKEDGNILGHKLEYNYNSHKVYITADKQKNGIKVNFKTRHNIEDGSVQLADHYQQNT<br>PIGDGPVLLPDNHYLSTQSALSKDPNEKRDHMVLLEFVTAAGITLGMDELYK* | 41320.2 | mEmerald-H2B-6<br>Addgene: 54111 |
| H2B-MiniR-mEmerald | MPEPAKSAPAPKKGSKKAVTKAQKKGGKKRKRSRKESYSIYVYKVLKQVHPDTGISSKAMGIM<br>NSFVNDIFERIAGEASRLAHYNKRSTITSREIQTAVRLLLPGELAKHAVSEGTKAITKYTSAKDP<br>M**LEIRVAFLRQRNTALRTEVAELEQEVQRLENRYGPL**PVATMVSKGEELFTGVVPILVELDGD<br>VNGHKFSVSGEGEGDATYGKLTLKFICTTGKLPVPWPTLVTTLTYGVQCFARYPDHMKQHDF<br>FKSAMPEGYVQERTIFFKDDGNYKTRAEVKFEGDTLVNRIELKGIDFKEDGNILGHKLEYNYNS<br>HKVYITADKQKNGIKVNFKTRHNIEDGSVQLADHYQQNTPIGDGPVLLPDNHYLSTQSKLSKDP<br>NEKRDHMVLLEFVTAAGITLGMDELYK* | 45802.4 | H2B-MiniR-mEm<br>Addgene:181990 |
| H2B-MiniR | MPEPAKSAPAPKKGSKKAVTKAQKKGGKKRKRSRKESYSIYVYKVLKQVHPDTGISSKAMGIM<br>NSFVNDIFERIAGEASRLAHYNKRSTITSREIQTAVRLLLPGELAKHAVSEGTKAITKYTSAKDP<br>M**LEIRVAFLRQRNTALRTEVAELEQEVQRLENRYGPL**LPVAT* | 18865.9 | H2B-MiniR |
| mCherry-TOMM20 | MVGRNSAIAAGVCGALFIGYCIYFDRKRRSDPNFKNRLRERRKKQKLAKERAGLSKLPDLKDA<br>EAVQKFFLEEIQLGEELLAQGEYEKGVDHLTNAIAVCGQPQQLLQVLQQTLPPPVFQMLLTKLP<br>TISQRIVSAQSLAEDDVEGGSGDPPVAT**MVSKGEEDNMAIIKEFMRFKVHMEGSVNGHEFEIE**<br>**GEGEGRPYEGTQTAKLKVTKGGPLPFAWDILSPQFMYGSKAYVKHPADIPDYLKLSFPEGF**<br>**KWERVMNFEDGGVVTVTQDSSLQDGEFIYKVKLRGTNFPSDGPVMQKKTMGWEASSERMY**<br>**PEDGALKGEIKQRLKLKDGGHYDAEVKTTYKAKKPVQLPGAYNVNIKLDITSHNEDYTIVEQY**<br>**ERAEGRHSTGGMDELYK*** | 43840.9 | mCherry-TOMM20-N-10<br>Addgene:55146 |
| TOMM20-PuRRRE | MVGRNSAIAAGVCGALFIGYCIYFDRKRRSDPNFKNRLRERRKKQKLAKERAGLSKLPDLKDA<br>EAVQKFFLEEIQLGEELLAQGEYEKGVDHLTNAIAVCGQPQQLLQVLQQTLPPPVFQMLLTKLP<br>TISQRIVSAQSLAEDDVEGGSGDPPVAT**LEIRNAFLRQRVTALRTRVSELRQENSELENEYGP**<br>**L*** | 21420.6 | TOMM20-PuRRRE |
| mEmerald-Actin | MVSKGEELFTGVVPILVELDGDVNGHKFSVSGEGEGDATYGKLTLKFICTTGKLPVPWPTLVTT<br>LTYGVQCFARYPDHMKQHDFFKSAMPEGYVQERTIFFKDDGNYKTRAEVKFEGDTLVNRIELK<br>GIDFKEDGNILGHKLEYNYNSHKVYITADKQKNGIKVNFKTRHNIEDGSVQLADHYQQNTPIGD<br>GPVLLPDNHYLSTQSKLSKDPNEKRDHMVLLEFVTAAGITLGMDELYKSGLRSGSGGGGSASG<br>GSGSDDDIAALVVDNGSGMCKAGFAGDDAPRAVFPSIVGRPRHQGVMVGMGQKDSYVGDE<br>AQSKRGILTLKYPIEHGIVTNWDDMEKIWHHTFYNELRVAPEEHPVLLTEAPLNPKANREKMTQ<br>IMFETFNTPAMYVAIQAVLSLYASGRTTGIVMDSGDGVTHTVPIYEGYALPHAILRLDLAGRDLT<br>DYLMKILTERGYSFTTTAEREIVRDIKEKLCYVALDFEQEMATAASSSSLEKSYELPDGQVITIG<br>NERFRCPEALFQPSFLGMESCGIHETTFNSIMKCDVDIRKDLYANTVLSGGTTMYPGIADRMQ<br>KEITALAPSTMKIKIIAPPERKYSVWIGGSILASLSTFQQMWISKQEYDESGPSIVHRKCF* | 69948.4 | mEmerald-Actin-C-18<br>Addgene:53978 |
| mEmerald-TREER-Actin | MVSKGEELFTGVVPILVELDGDVNGHKFSVSGEGEGDATYGKLTLKFICTTGKLPVPWPTLVTT<br>LTYGVQCFARYPDHMKQHDFFKSAMPEGYVQERTIFFKDDGNYKTRAEVKFEGDTLVNRIELK<br>GIDFKEDGNILGHKLEYNYNSHKVYITADKQKNGIKVNFKTRHNIEDGSVQLADHYQQNTPIGD<br>GPVLLPDNHYLSTQSKLSKDPNEKRDHMVLLEFVTAAGITLGMDELYKSGLRS**LEIRVAFLRQE**<br>**VTALETENAELEQRVQRLRNRYGPL**GGGRSGSGGGSASGGSGSDDDIAALVVDNGSGMCK<br>AGFAGDDAPRAVFPSIVGRPRHQGVMVGMGQKDSYVGDEAQSKRGILTLKYPIEHGIVTNWD<br>DMEKIWHHTFYNELRVAPEEHPVLLTEAPLNPKANREKMTQIMFETFNTPAMYVAIQAVLSLYA<br>SGRTTGIVMDSGDGVTHTVPIYEGYALPHAILRLDLAGRDLTDYLMKILTERGYSFTTTAEREIV<br>RDIKEKLCYVALDFEQEMATAASSSSLEKSYELPDGQVITIGNERFRCPEALFQPSFLGMESCG<br>IHETTFNSIMKCDVDIRKDLYANTVLSGGTTMYPGIADRMQKEITALAPSTMKIKIIAPPERKYSV<br>WIGGSILASLSTFQQMWISKQEYDESGPSIVHRKCF* | 74656.7 | mEmerald-TREER-Actin |
| TREER-Actin | M**LEIRVAFLRQEVTALETENAELEQRVQRLRNRYGPL**GGGRSGSGGGSASGGSGSDDDIAA<br>LVVDNGSGMCKAGFAGDDAPRAVFPSIVGRPRHQGVMVGMGQKDSYVGDEAQSKRGILTLK<br>YPIEHGIVTNWDDMEKIWHHTFYNELRVAPEEHPVLLTEAPLNPKANREKMTQIMFETFNTPA<br>MYVAIQAVLSLYASGRTTGIVMDSGDGVTHTVPIYEGYALPHAILRLDLAGRDLTDYLMKILTER<br>GYSFTTTAEREIVRDIKEKLCYVALDFEQEMATAASSSSLEKSYELPDGQVITIGNERFRCPEAL<br>FQPSFLGMESCGIHETTFNSIMKCDVDIRKDLYANTVLSGGTTMYPGIADRMQKEITALAPSTM<br>KIKIIAPPERKYSVWIGGSILASLSTFQQMWISKQEYDESGPSIVHRKCF* | 49442.3 | TREER-Actin |

**Table S2: Bacterial strains and plasmids.**

| *E. coli* Strains | Characteristics | Source |
|---|---|---|
| Top10 | F- mcrA Δ(mrr-hsdRMS-mcrBC) Φ80lacZΔM15 Δ lacX74 recA1 araD139 Δ(araleu)7697 galU galK rpsL (StrR) endA1 nupG | Thermo Fisher Scientific |
| BL21(DE3) | F- ompT hsdSB (rBmB-) gal dcm (DE3) | Thermo Fisher Scientific |
| NEB-5alpha | fhuA2Δ(argF-lacZ)U169 phoA glnV44 Φ80Δ(lacZ)M15 gyrA96 recA1 relA1 endA1 thi-1 hsdR17 | New England Biolabs |
| **Plasmids** | **Characteristics** | **Source** |
| pET28b (+) | T7 promoter, His-tag coding sequence, MCS, *lacI* coding sequence, (KanR) | Novagen |
| pcDNA3.1 | CMV promoter, MCS, BGH polyadenylation signal, SV40 origin, (AmpR, KanR) | Thermo Fisher Sci |
| mCherry-TOMM20-N-10 | CMV promoter, TOMM20, mCherry (C terminal on backbone), (KanR, NeoR) | Addgene: 55146 |
| H2B-6-mEmerald | CMV promoter, HIST1H2BJ, mEmerald (C terminal on backbone), (KanR,NeoR) | Addgene: 54111 |
| mEmerald-Actin-C-18 | CMV promoter, actin, mEmerald (N terminal on the backbone), (KanR, NeoR) | Addgene: 53978 |

**Table S3: Properties of fluorophores.**

| Fluorophore | Vendor | Excitation Maximum | Emission Maximum | Quantum Yield | Extinction Coefficient (ε) | Correction $Factor_{280}$ |
|---|---|---|---|---|---|---|
| Sulfo-Cyanine5-Maleimide (Sulfo-Cy5) | Lumiprobe | 646 nm | 662 nm | 0.28 | 271,000 | 0.04 |
| Sulfo-Cyanine3-Maleimide (Sulfo-Cy3) | Lumiprobe | 548 nm | 563 nm | 0.1 | 162,000 | 0.06 |
| AlexaFluor 488 $C_5$ Maleimide (AF488) | Thermo Fisher Sci | 493 nm | 516 nm | 0.92 | 72,000 | 0.11 |

**Table S4: Summary of probe peptides.**

| Peptide | Fluorophore | Conc. (μM) | % labeled | Lot |
|---|---|---|---|---|
| MiniE-Cy5 | Sulfo-Cy5 | 43 | 75 | KD20210420 |
| MiniE-Cy3 | Sulfo-Cy3 | 8.5 | 90 | KD20210608 |
| MiniE-AF488 | AF488 | 6.8 | 100 | KD20220923 |
| MiniR-Cy5 | Sulfo-Cy5 | 35 | 29 | JD20190715 |
| TinyERRE-Cy5 | Sulfo-Cy5 | 102 | 54 | KD20220107 |
| TinyERRE-Cy3 | Sulfo-Cy3 | 8 | 50 | AS20210607 |
| PuRRRE-Cy5 | Sulfo-Cy5 | 4.6 | 85 | AS20210614 |
| PunyEEER-Cy5 | Sulfo-Cy5 | 14 | 54 | AS20210614 |

*Attempts to label TREER never produced a viable probe peptide labeled greater than 20%.

### References


(1) Doh, J. K.; Tobin, S. J.; Beatty, K. E. MiniVIPER Is a Peptide Tag for Imaging and Translocating Proteins in Cells. *Biochemistry* **2020**, *59* (33), 3051-3059. DOI: 10.1021/acs.biochem.0c00526.

(2) Zane, H. K.; Doh, J. K.; Enns, C. A.; Beatty, K. E. Versatile Interacting Peptide (VIP) Tags for Labeling Proteins with Bright Chemical Reporters. *ChemBioChem* **2017**, *18* (5), 470-474. DOI: 10.1002/cbic.201600627.

(3) Sambrook, J.; Russell, D. W. Purification of Expressed Proteins from Inclusion Bodies. *Cold Spring Harbor Protocols* **2006**, *2006* (1), pdb.prot4089. DOI: 10.1101/pdb.prot4089.

(4) *The QIAexpressionist*; Qiagen, 2003.

(5) Moll, J. R.; Ruvinov, S. B.; Pastan, I.; Vinson, C. Designed heterodimerizing leucine zippers with a ranger of pIs and stabilities up to 10−15 M. *Protein Science* **2001**, *10* (3), 649-655. DOI: https://doi.org/10.1110/ps.39401.

(6) Litowski, J. R.; Hodges, R. S. Designing heterodimeric two-stranded alpha-helical coiled-coils. Effects of hydrophobicity and alpha-helical propensity on protein folding, stability, and specificity. *J Biol Chem* **2002**, *277* (40), 37272-37279. DOI: 10.1074/jbc.M204257200

(7) Kantner, T.; Alkhawaja, B.; Watts, A. G. In Situ Quenching of Trialkylphosphine Reducing Agents Using Water-Soluble PEG-Azides Improves Maleimide Conjugation to Proteins. *ACS Omega* **2017**, *2* (9), 5785-5791. DOI: 10.1021/acsomega.7b01094.

(8) McGraw, T. E.; Greenfield, L.; Maxfield, F. R. Functional expression of the human transferrin receptor cDNA in Chinese hamster ovary cells deficient in endogenous transferrin receptor. *J Cell Biol* **1987**, *105* (1), 207-214. DOI: 10.1083/jcb.105.1.207

(9) Schindelin, J.; Arganda-Carreras, I.; Frise, E.; Kaynig, V.; Longair, M.; Pietzsch, T.; Preibisch, S.; Rueden, C.; Saalfeld, S.; Schmid, B.; et al. Fiji: an open-source platform for biological-image analysis. *Nat Methods* **2012**, *9* (7), 676-682. DOI: 10.1038/nmeth.2019

(10) Pearson, K. Mathematical contributions to the theory of evolution III. Regression, heredity and panmixia. *Philos Trans R Soc Lond B Biol Sci* **1896**, (187), 253-318. Dunn, K. W.; Kamocka, M. M.; McDonald, J. H. A practical guide to evaluating colocalization in biological microscopy. *Am J Physiol Cell Physiol* **2011**, *300* (4), C723-742. DOI: 10.1152/ajpcell.00462.2010
